# Structural basis of DegP-protease temperature-dependent activation

**DOI:** 10.1101/2020.07.14.202507

**Authors:** Darius Sulskis, Johannes Thoma, Björn M. Burmann

## Abstract

Protein quality control is an essential cellular function mainly executed by a vast array of different proteases and molecular chaperones. One of the bacterial HtrA (high temperature requirement A) protein family members, the homo-oligomeric DegP-protease, plays a crucial role in the *Escherichia coli* (*E. coli*) protein quality control machinery by removing unfolded proteins or preventing their aggregation and chaperoning them to their final folded state within the periplasm. DegP contains two regulatory PDZ domains, which play key roles in substrate recognition and in the transformation of DegP between inactive hexameric and proteolytic active cage-like structures. Here, we analyze the interaction and dynamics of the DegP PDZ-domains underlying this transformation by high-resolution NMR spectroscopy complemented with biochemical cleavage assays. We identify an interdomain molecular lock, which controls the interactions between the two PDZ domains, regulated by fine-tuned temperature-dependent protein dynamics, and which is potentially conserved in proteins harboring tandem PDZ domains.

## Introduction

An effective protein quality control (PQC) system consisting of molecular chaperones as well as proteases is essential for each living organism (*1, 2*). Within the periplasmic space of Gram-negative bacteria, this crucial task is carried out mainly by the following three proteins: Skp, SurA, and DegP (*3*). These proteins play a pivotal role in the quality control of outer membrane proteins (Omp) and protect bacteria from the accumulation and aggregation of misfolded proteins within the periplasm. Skp and SurA are responsible for controlling protein folding through their function as holdase chaperones (*4, 5*). In contrast, DegP has been shown to have a tightly-regulated dual activity under different stress conditions *in vivo* (*3*): Primarily, it functions as a serine protease to rescue cells during heat stress, but it also possesses chaperoning activity at reduced temperatures (*6*). DegP was shown to be essential for *E. coli* survival as a *degP* gene knockout is lethal at elevated temperatures (42°C) (*7*). However, it was possible to show that DegP protease activity is not a mandatory feature since overexpression of protease-deficient forms of DegP is already sufficient for *E. coli* to recover (*8*) under stress conditions, suppressing the lethal phenotypes (*9*).

On a structural level, DegP consists of a protease domain connected to two adjunct PDZ domains named PDZ1 and PDZ2 after the first proteins PSD-95, Dlg1, and ZO-1 in which this type of domains was observed (*10*). In its protease-inactive state DegP forms a homo-hexameric complex (*11*). Upon activation DegP can re-arrange into dodecameric or 24–meric cage-like forms, which are suggested to encapsulate substrate proteins before proteolytic cleavage (*12–14*) (Fig. 1A, B). The common building block of the different cage-like structures is a trimer, therefore DegP is assumed to undergo large structural transitions from its hexameric resting state *via* a trimeric intermediate state towards the higher oligomeric states (*15, 16*). This structural transition itself is supposed to be mediated by modulations of intermolecular PDZ1– PDZ2 interactions (*17*). Moreover, several loops, termed LA, LD, L1, L2, and L3, play an important role in the regulation of the proteolytic function of DegP, having either activating or inhibitory roles (*12, 18*) (fig. S1A, B). Among these, the so-called LA-loop plays an important role as it is believed to stabilize the inactive conformation of DegP by shielding the active site within the protease domain in cellular ground states. In addition, it crucially participates in the modulation of DegP function as mutations in this region directly affect the activation and functionality of the enzyme (*18, 19*). Upon activation, the LA-loop is supposed to facilitate local rearrangements leading to the correct orientation of the catalytic triad ensuring effective proteolysis (*18, 19*).

**Fig. 1:**
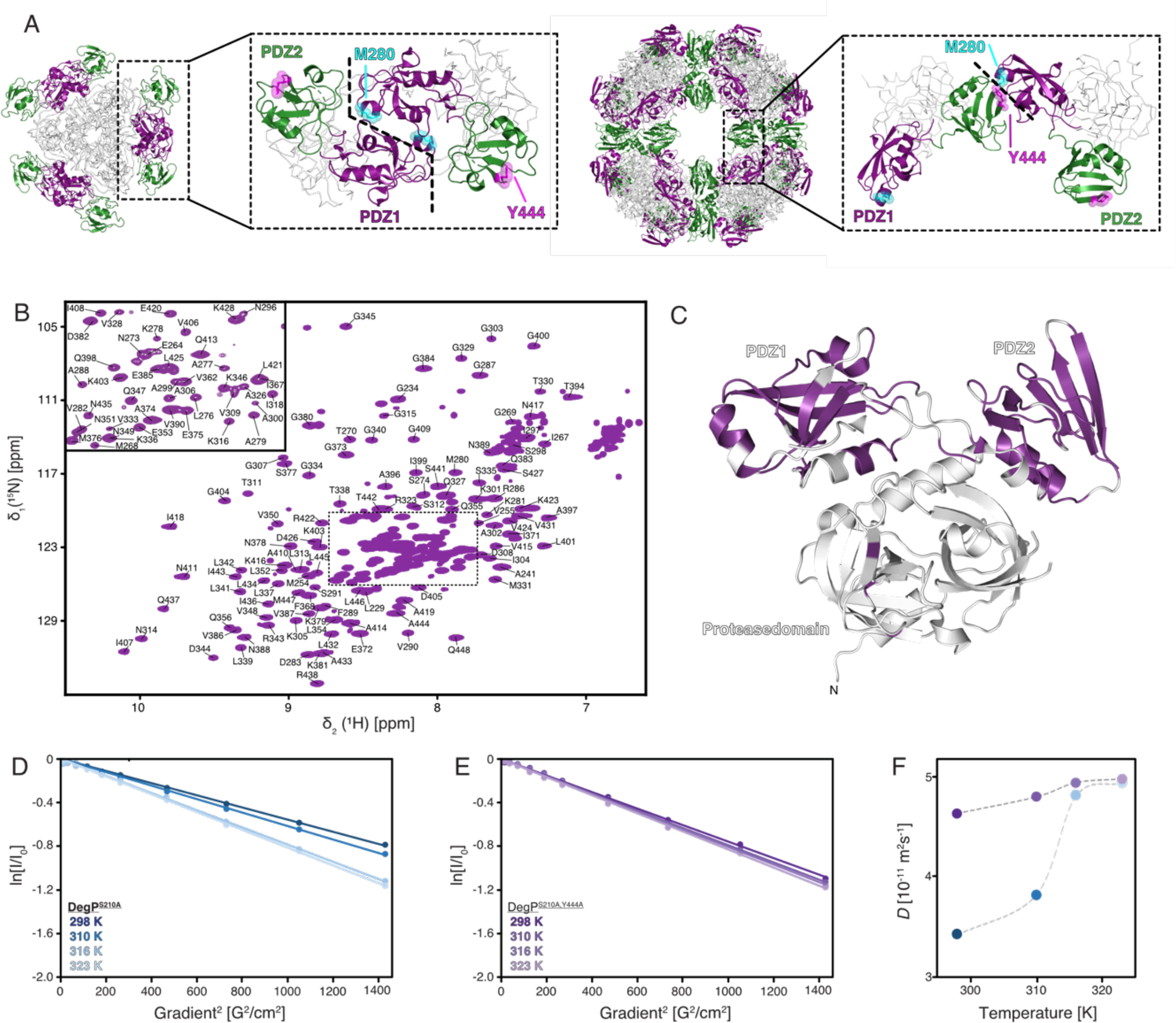
Full-length DegP^S210A^ in solution. (**A**) Comparison of the PDZ domains orientation between different monomers in the crystal structures of the hexamer (PDB-ID: 1KY9) and 24–meric cage assembly (PDB-ID: 3OU0). Whereas in the inactive hexameric state this interaction is proposed to be mediated mainly *via* a direct interaction between two PDZ1s, the PDZ2s are found to be non-constraint at the hexamer edges, however within the proteolytically active 24-mer the interaction between PDZ1 and PDZ2 of different molecules is achieved by a Y444 and M280 interaction indicating stabilization by a sulfur-*π*-aromatic or methyl-*π*-aromatic motif. (**B**) 2D [^15^N,^1^H]-NMR spectrum of [*U*-^2^H,^15^N,^13^C]-DegP^S210A,Y444A^. The sequence-specific resonance assignment obtained from 3D TROSY-type triple-resonance experiments is indicated. (**C**) Cartoon representation of a DegP^S210A^ monomer from the crystal structure of the hexameric assembly of DegP^S210A^ (PDB-ID: 1KY9) with the sequence-specific resonance assignment indicated in purple. (**D, E**) Measurement of the molecular diffusion constant with a ^13^C-methyl-filtered diffusion experiment for DegP^S210A^ (**D**) and DegP^S210A,Y444A^ (**E**). The logarithm of the signal intensity is plotted against the squared gradient strength of the applied pulsed-field gradients. The two [*U*-^2^H, Ile-*δ*_1_-^13^CH_3_]–DegP samples were measured at different temperatures as indicated. Solid lines are a linear fit to the data. (**F**) Obtained molecular diffusion constants plotted against the temperature. The broken lines serve as a guide to the eyes only.

Nevertheless, the exact underlying mechanism and eventual structural adaptions remain only partially understood as previous structural studies could only reveal the respective endpoints of the different oligomeric assemblies. The current model suggests that peptides containing a hydrophobic carboxy-terminal tail initially bind to the PDZ1-domain, thereby triggering subsequent structural re-arrangements, which in turn activate DegP (*20*). Contrary, the PDZ2-domain inhibits protease activity as long as DegP disassembles into trimers and subsequently transforms into the cage-like structures (*17*).

To address the underlying structural adaptions upon temperature activation of DegP, we investigated full-length DegP as well as its isolated PDZ1–PDZ2 domains at the atomic level using high-resolution NMR spectroscopy. Based on local chemical shift perturbations we identified an intermolecular methionine–tyrosine interaction controlling the transition from hexameric to trimeric DegP. Detailed analysis of the inherent protein dynamics revealed a finely tuned temperature-dependent allosteric network within PDZ1 modulated by the relief of the interaction with PDZ2. Biochemical characterization of the effect of this interdomain lock on the proteolytic function identified this temperature sensor as the first step in the activation cascade of DegP, but not sufficient for complete protease activation. Remarkably, the crucial tyrosine–methyl bearing residues are conserved within HtrA proteins, in perfect agreement with the mechanism of inhibitory transient oligomerization recently identified for human HtrA2 (*21*), indicating a widespread fundamental activation mechanism modulated *via* an intermolecular lock between the PDZ domains.

## Results

### Solution NMR of full-length DegP

Currently, only static high-resolution structures of DegP by X-crystallography (*13*) and medium-resolution structures by cryo-electron microscopy (cryo-EM) (*22*) are available, providing an incomplete picture of the functional cycle of DegP. While these structures provide important insight into the organization of the different oligomeric states of DegP, the underlying structural transitions could not be deduced from them. Therefore, we investigated DegP by high-resolution NMR-spectroscopy to obtain detailed insight into its structural and dynamical properties, which give rise to the different oligomeric states and coordinate the transition from the inactive to the active states of DegP. Due to its large size (∼150 kDa as a trimeric building block) and the existence of different oligomeric forms (Fig. 1A, B), DegP presents a challenging system for solution NMR studies. Here, we present the first [^15^N,^1^H]-NMR spectra of uniformly-labeled [*U*-^2^H,^15^N]-DegP^S210A^, which has the active site serine mutated to prevent self-proteolysis (sequence numbering follows the mature protein 1–448 excluding the 26 residues long periplasmic targeting sequence) (fig. S1C–E). The spectrum obtained at 25°C contained only a very limited number of resonances located exclusively in the central region of the spectrum, indicative of disordered and unstructured regions such as loops, in line with the large size of the protein (fig. S1C). In contrast, increasing the temperature to 37°C and subsequently, to 50°C resulted in well-dispersed spectra containing approximately 120 resonances (fig. S1D, E). The dispersion of these peaks [10–6 ppm in the ^1^H-dimension] showed that a folded domain of the full-length DegP was observable under these conditions, indicating its rather flexible attachment to the rest of the protein. In the next step, we used the known trimerization mutant DegP^S210A,Y444A^ (*23*), which resulted in further spectral improvement (Fig. 1C and fig. S1F) and allowed us to sequence-specifically assign large parts of the two carboxy-terminal PDZ domains (150 out of a total of 188 PDZ1–PDZ2 resonances) using standard TROSY (transverse relaxation-optimized spectroscopy)-type 3D experiments. The obtained secondary chemical shifts indicated that the structure in solution is to a large extent in agreement with the secondary structure elements observed previously within the different crystal structures (fig. S1E–G). The observation of a large set of resonances of the DegP PDZ domains indicates that these domains are not in direct contact with other parts of the protein and tumble independently in solution. Furthermore, the increased spectral quality at elevated temperatures suggested that full-length DegP might transition between different oligomeric states, pointing to an inherent temperature switch for DegP activation.

In order to determine the oligomeric state of full-length DegP^S210A^ in solution, we initially tried to use the TRACT experiment (*24*) to probe the DegP size at the elevated temperatures (data not shown). However, due to the lack of signals at lower temperatures, this approach could not be used at temperatures below 37°C. Moreover, the inherent overestimation of flexible regions in the TRACT experiment prevented in-depth analysis (*24*). To circumvent these issues, we specifically labeled the methyl groups of the methyl bearing isoleucine residues, which yielded high-quality spectra already at 25°C, and measured pulsed-field gradient diffusion experiments on the methyl groups (*25*). The obtained translational diffusion coefficients showed a clear temperature dependence (Fig. 1D). With increasing temperature, the values increased from 3.4•10^-7^ cm^2^s^-1^ at 25°C to 4.82•10^-7^ cm^2^s^-1^ and 4.95•10^-7^ cm^2^s^-1^ at 43°C and 50°C, respectively (Fig. 1D–F). Comparing these values to known literature values clearly indicates that DegP is in a hexameric state (MW = 300 kDa) at 25°C whereas at 43°C and higher the protein is mainly in a trimeric state (MW = 150 kDa) (fig. S1J). At 37°C we obtained diffusion coefficients that were slightly elevated compared to 25°C (Fig. 1D–F) indicating the partial presence of a trimeric species as an initial step of the structural transition, which is estimated to be about 20– 25% based on the increase in the diffusion coefficient. Our observation of a temperature dependent disassembly of the DegP hexamer is in complete agreement with earlier size exclusion studies indicating the formation of trimeric DegP^S201A^ (∼80 % populated) at heat shock temperatures of 42°C (*17*). To further verify this observation we probed the effects of the point mutation DegP^S210A,Y444A^. In contrast to DegP^S210A^ this variant eluted from a size-exclusion column as a mixture of trimer and hexamer already at 8°C (fig. S1D). To address the temperature dependent modulation of the oligomeric states for this variant, we also employed NMR diffusion measurements. With increasing temperature, the values only showed a slight variation, increasing from 4.64•10^-7^ cm^2^s^-1^ at 25°C to 4.81•10^-7^ cm^2^s^-1^ and 4.99•10^-7^ cm^2^s^-1^ at 37°C and 50°C, respectively (Fig. 1E, F and table S1). In comparison to DegP^S210A^, these values indicate that already at 25°C the majority of this variant is in the trimeric state. Importantly, the observation of trimerization upon mutating Y444 to alanine and the possible temperature dependent release of the PDZ domains in wild-type DegP cannot be directly explained by the known hexameric DegP structure (Fig. 1A) (*13*). In fact, within the larger oligomeric states, a domain lock is observed on the interface formed by Y444 and M280 (Fig. 1B) (*22*). To address whether this Y444–M280 interaction occurred as well in solution within the hexamer to trimer transition, we also tested the effect of mutating the corresponding methionine. Size-exclusion chromatography revealed the existence of distinct trimeric and hexameric species for the DegP^S210A,M280A^ variant, in line with the hypothesis of an Y444– M280 interaction modulating the transition of the oligomeric states (Fig. 1D). To gain additional insight into the nature of the interaction between these two crucial residues we additionally tested the more conservative mutation variant DegP^S210A,M280I^. In contrast, this variant showed no distinct states in the size exclusion chromatography, but rather a single elution peak in between the trimeric and the hexameric state, indicating a fast trimer-hexamer equilibrium suggesting a weakened complex formation compared the DegP^S210A^, which forms an obligate hexamer under the tested conditions (fig. S1F). In agreement with this observation, we obtained also a slightly altered hexamer-trimer transition when studying this DegP^S210A,M280I^ variant by NMR-diffusion (fig. S1H-I).

### Isolated PDZ1–PDZ2 bi-domains in solution

Based on the initial observation of the highly flexible PDZ-domains within the full-length DegP and their possible contribution to the inherent temperature switch, we subsequently focused on the isolated PDZ1–PDZ2 domains. High-quality resonance assignment data of the PDZ1–PDZ2 domain construct allowed an almost complete sequence-specific assignment of the observed resonances in the [^15^N,^1^H]-NMR spectrum (fig. S2), yielding 87% backbone and 95% aliphatic side-chain assignment. The missing amino acids were part of the interdomain linker (356–366) or were located in flexible loops within PDZ1. This observed exchange broadening can be explained by enhanced amide-proton exchange rates at 50°C within these unstructured regions (*26*). Using the combined ^13^C*α*and ^13^C*β* chemical shifts of the protein backbone, we identified the secondary structure elements of the PDZ1–PDZ2 domain construct in solution. The observed structural elements, five *α*-helices and ten *β*-sheets, are in agreement with the apparent secondary structure elements of the available crystal structures (fig. S2C, D). However, we observed some difference within PDZ2, where the short helical turn *α*_4_ was slightly shifted and the *β*-strand *β*_8_ also lacked the characteristic secondary chemical shifts. Nevertheless, the presence of the short *β*-strands, *β*_6_ and *β*_7_, in the linker region between the two PDZ domains, suggests that the domains are not completely decoupled, even at the elevated temperature of 50°C.

As we had observed large temperature-dependent effects within the spectra of full-length DegP, we also assigned the PDZ1–PDZ2 construct at 25°C and investigated its secondary structure propensities (fig. S3). The ^13^C secondary chemical shifts were almost identical to the ones obtained at 50°C, indicating that no temperature-dependent structural adaptions occurred within the PDZ domains. Nevertheless, a distinct lack of resonances was observed for residues within the amino- and carboxy-terminus of the PDZ1–PDZ2 domain construct in the 25°C data set: the resonances for residues M268, G269, L276, A277, M280, R286, G287, S291, N435, I436, G439, D440, and I443–M447 were broadened beyond detection in a [^15^N,^1^H]-NMR spectrum, in line with the presence of a dynamic intermolecular interaction between different PDZ1–PDZ2 molecules on the NMR intermediate timescale, i.e. with kinetic rates ranging from 1000 to 10 s^-1^ (*27*).

### Probing the interactions between PDZ1 and PDZ2

To follow up on the temperature-dependent line-broadening effect observed for the PDZ1– PDZ2 domain construct in solution, we characterized the PDZ1–PDZ2 interaction in further detail. Previous cryo-EM data indicated that the PDZ1–PDZ2 domains are essential for assembling as well as interlocking the DegP 12–meric and 24–meric cages and that removal of the PDZ2 domain prevents cage formation, altogether leading to primarily trimeric protein (*17*). Therefore, we prepared individual constructs of the PDZ1 and the PDZ2 domain and confirmed the sequence specific resonance assignments (fig. S4). In order to map possible interaction areas between these two isolated domains at different temperatures, we titrated the PDZ1 and the PDZ2 domains towards each other at four different temperatures (25–50°C).

These temperatures represent the hexameric, the ∼20% trimeric, as well as the fully trimeric state within the full-length DegP^S210A^ protein, as indicated by the NMR diffusion data (Fig. 1D–F). The observed spectral changes showed that the extreme amino-terminal and carboxy-terminal regions of the PDZ1–PDZ2 domains interact strongly at 25°C and the chemical shift perturbations (CSPs) lessened with increasing temperature, with M280 showing the largest effect (Fig. 2 and fig. S5). Remarkably, completely in line with our previous observations for the larger DegP constructs, the affected resonances did not correspond to any interfaces on the available hexameric crystal structure (Fig. 1A), but were observed within the 12 meric X-ray and the 24-meric cryo-EM structure (Fig. 1B and 2E), indicating the possibility of a stabilizing sulfur-π aromatic or methyl-π aromatic motif (*28, 29*). Moreover, a distinct set of residues showed larger chemical shift perturbations along with an enhanced intensity loss at 25°C compared to higher temperatures (Fig. 2C, D). Specifically, besides M280 Y444 was the residue most affected, implying that these two residues might provide the molecular switch locking the hexameric resting state at lower temperatures, completely in line with our observations by size-exclusion chromatography and NMR diffusion for the different full-length protein variants (Fig. 1D–F). To validate this hypothesis, we constructed the corresponding mutants (PDZ1^M280A^, PDZ2^Y444A^) and evaluated if the interaction between the domains was abolished. The titration experiments clearly showed that the interaction was completely abolished if either M280 or Y444 was mutated (fig. S6A, B), highlighting the involvement of both amino acids in forming the intermolecular lock. The same characteristic effect, although to a slightly lesser extent, was observed for the sub-set of peaks involved in the interaction for the PDZ^M280I^ variant, indicating that the complex was less stable (fig. S6C).

**Fig. 2:**
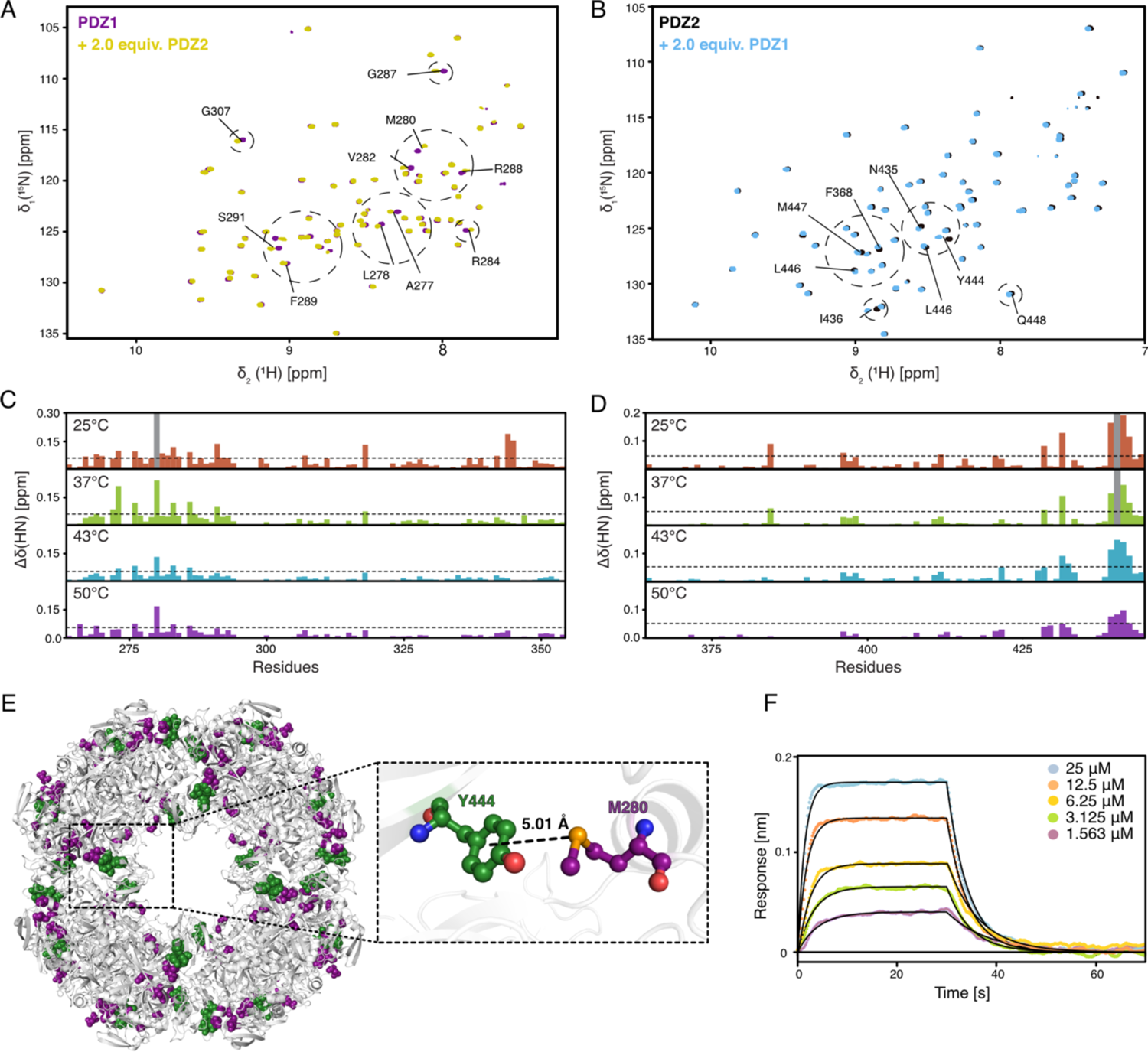
The individual PDZ domains show transient interactions reminiscent of the higher oligomeric DegP-complexes. (**A, B**) 2D [^15^N,^1^H]-NMR spectra of [*U*-^15^N]-PDZ1 (**A**, purple) and [*U*-^15^N]-PDZ2 (**B**, black) as well as after the addition of two molar equivalents of the respective other domain. Titrations were measured at 50°C. (**C, D**) Detected chemical shift perturbations at four temperatures ranging from 25°C–50°C as indicated by the PDZ1 domain (**C**) and the PDZ2 domain (**D**). Severely affected residues experiencing line-broadening are indicated as grey bars in the panels. The dotted lines represent a significance level of 0.05 ppm of the CSPs. (**E**) Perturbated resonances mapped on the crystal structure of DegP^S210A^ 24–mer (PDB-ID: 3OU0). Mainly affected residues, CSPs twice the S.D. on PDZ1, are highlighted in purple (G266, G269, K278, M280, D283, and R286), whereas residues in PDZ2 are highlighted in green (A433, N435, I443, Y444, and L445). The enlargement shows the central M280–Y444 interaction. (**F**), Kinetic analysis by biolayer interferometry (BLI) of the PDZ1– PDZ2 interaction. PDZ1 binding to the biotinylated-PDZ2 domain was probed at 25°C. Analyte concentrations are indicated in the figure. Non-linear least-square fits to the experimental data are indicated by black lines.

To quantify the interaction between the individual wild-type PDZ domains we used Bio-Layer Interferometry (BLI) steady-state analysis at 25°C resulting in a dissociation constant (*K*_D_) of 7.8 ± 0.9 µM for the binding of PDZ1 towards biotinylated-PDZ2 (Fig. 2F and fig. S4C). We did not test the interaction *vice versa* with biotinylated PDZ1 in detail, because initial tests yielded a dissociation constant weakened by a factor of ∼20, indicating that the binding interface on PDZ1 is perturbed by the close proximity of the Biotin-label and the identified interaction site (data not shown).

### The M280–Y444 interaction locks the hexameric state

To further analyze the basis of the M280–Y444 interaction we initially checked the temperature effects on the resonance lines of the side-chains within the PDZ1–PDZ2 construct. For Y444 we observed a C*α* resonance line-broadening to a similar extent than observed initially for the amide moiety (Figs. 2B and 3A). This behavior is furthermore reflected by the Y444*δ* ring resonances, which lose about 60% of their intensity at 25°C compared to 50°C in contrast to the Y444*ε* resonances, which show stable intensity over the tested temperature range (Fig. 3B). Subsequently, we investigated the effects on the M280 side, where we focused on its methionine-*ε*-methyl group, which we assigned by employing the mutation variant PDZ1– PDZ2^M280A^ (Fig. 3C). Probing the M280*ε* resonance at different temperatures revealed the presence of two distinct states, which we termed open (50°C) and locked (25°C) (Fig. 3D). In the transition temperatures of 37°C and 43°C, respectively, we could observe two distinct peaks for the open and locked conformations, which are in the slow exchange regime, milliseconds to seconds, on the NMR timescale. Integrating the peak intensities at the different temperatures revealed that at 37°C the open and locked conformation coexist in a 1:1 ratio, whereas at 43°C or higher the open conformation dominates, and at 25°C the locked one, respectively (Fig. 3E). In order to quantify the interconversion between the open and the locked state, we used NMR exchange spectroscopy (Fig. 3F,G) (*31, 32*). We estimated the rate for the interconversion between the different states of *k*_ex_ = 0.33 s^-1^ from the open to the locked state, although this has to be considered as a rough estimate as the overlap of the second cross peak impaired a more detailed analysis. Interestingly, the observation of an opening-mechanism on this slow exchange timescale is in line with recently reported data on a trimer–hexamer equilibrium for the related human HtrA2 (*21*).

**Fig. 3:**
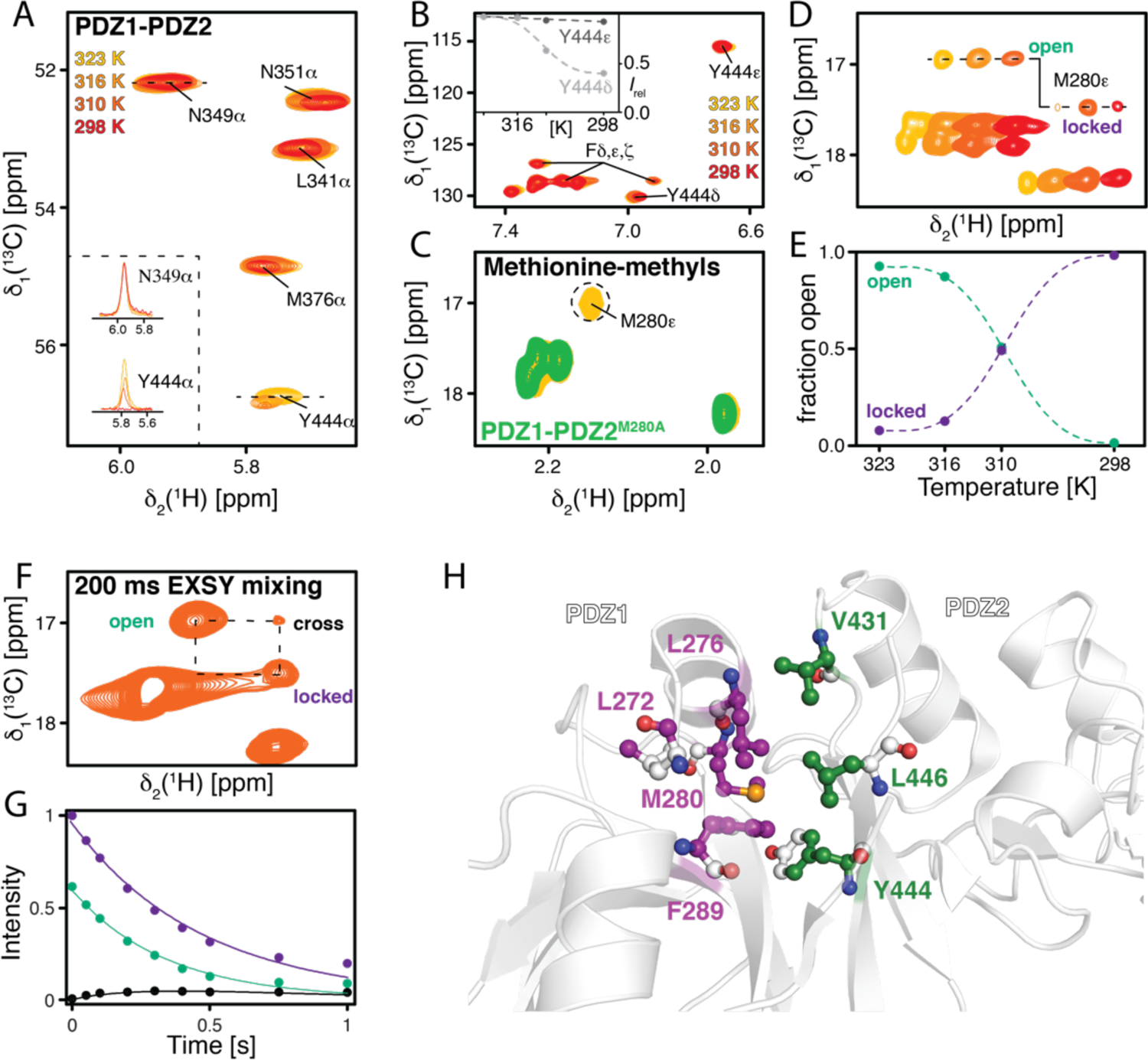
M280 is locking the PDZ1–PDZ2 interaction within DegP *via* Y444. (**A**) Focus on the C*α* resonance region around Y444 of a 2D [^13^C,^1^H]-NMR spectrum of [*U*-^13^C,^15^N]-PDZ1–PDZ2 at 25°C– 50°C as indicated. The inset shows the respective *δ*_2_[^1^H]-1D cross–sections through the N349*α* and Y444*α* resonances. (**B**) Methionine methyl region of a 2D [^13^C,^1^H]-NMR spectrum of [*U*-^13^C,^15^N]-PDZ1–PDZ2 at 25°C–50°C as indicated. Inset shows the signal intensities of the aromatic Y444*ε*and Y444*δ* resonances in the dependence of the temperature. (**C**) Overlay of [*U*-^13^C,^15^N]-PDZ1–PDZ2^M280A^ (blue) and PDZ1–PDZ2 (yellow) at 50°C used to assign the M280*ε* resonance. (**D**) Methionine-*ε*-methyl region of a 2D [^13^C,^1^H]-NMR spectrum of [*U*-^13^C,^15^N]-PDZ1–PDZ2 at 25°C–50°C as indicated. Spectra were manually shifted along the ^1^H dimension to illustrate the transition between open and locked states. (**E**) Analysis of the open and locked state. Lines are guide to the eyes only. (**F**) Magnetization exchange spectroscopy showing that the open state (o) and locked state (l) of M280 are in a dynamic equilibrium at 37°C as indicated. (**G**) The exchange rate (*k*_ex_) is determined from the built-up of the exchange peaks. (**H**) Observed NMR line-broadening indicates a hydrophobic patch stabilizing the PDZ1–PDZ2 interaction. Atoms involved on PDZ2 side are highlighted in green, whereas residues on PDZ1 side are highlighted in purple.

To obtain a more detailed picture of the involved residues in the stabilization of this interdomain lock we analyzed nuclear Overhauser enhancement spectroscopy (NOESY) cross-peaks in a temperature-dependent manner (fig. S7). This analysis revealed that the participating residues, L276, M280, F289 within PDZ1 as well as V431, Y444, L446 within PDZ2 are involved in stable NOE networks in their respective domains over the temperature range 37°C– 50°C. Due to the line-broadening of the participating atoms no intermolecular NOEs could be detected at 25°C, but the distinctive signal intensity loss clearly showed the regions involvement in this temperature lock by forming a hydrophobic patch, in agreement with the interface identified in the structures of the larger DegP assemblies earlier (Fig. 2H) (*22*). Interestingly, we also observed temperature-dependent line-broadening for the methyl moieties of L272, which indicates its contribution to the interaction network. However, based on the crystal structure of the DegP_12_(*23*), this residue is not expected to be located directly within the binding patch, indicating a slight rearrangement of this part of the protein, potentially due to the isolated PDZ1–PDZ2 construct used.

### Backbone dynamics of the PDZ1–PDZ2 construct reveal a fine-tuned temperature adaption

In order to understand the molecular basis and eventual structural or functional consequences of the M280 mediated temperature-switch in more detail, we next evaluated the inherent backbone dynamics of the PDZ1–PDZ2 domain construct over a broad range of timescales employing NMR relaxation measurements (*32*). To investigate the motions on the fast-timescale (pico- to nanosecond) the steady-state ^15^N{^1^H}-NOE (hetNOE) and the ^15^N longitudinal (*R*_1_) relaxation rates were measured. We observed a planar profile for the folded segments with average hetNOE values of 0.78 and *R*_1_-rates of 1.82 s^-1^, respectively, indicating a stable protein fold (Fig. 4A and fig. S8A). As expected, for the loop regions and the termini we observed values indicative of increased flexibility, namely lower hetNOE values and higher *R*_1_-rates.

**Fig. 4:**
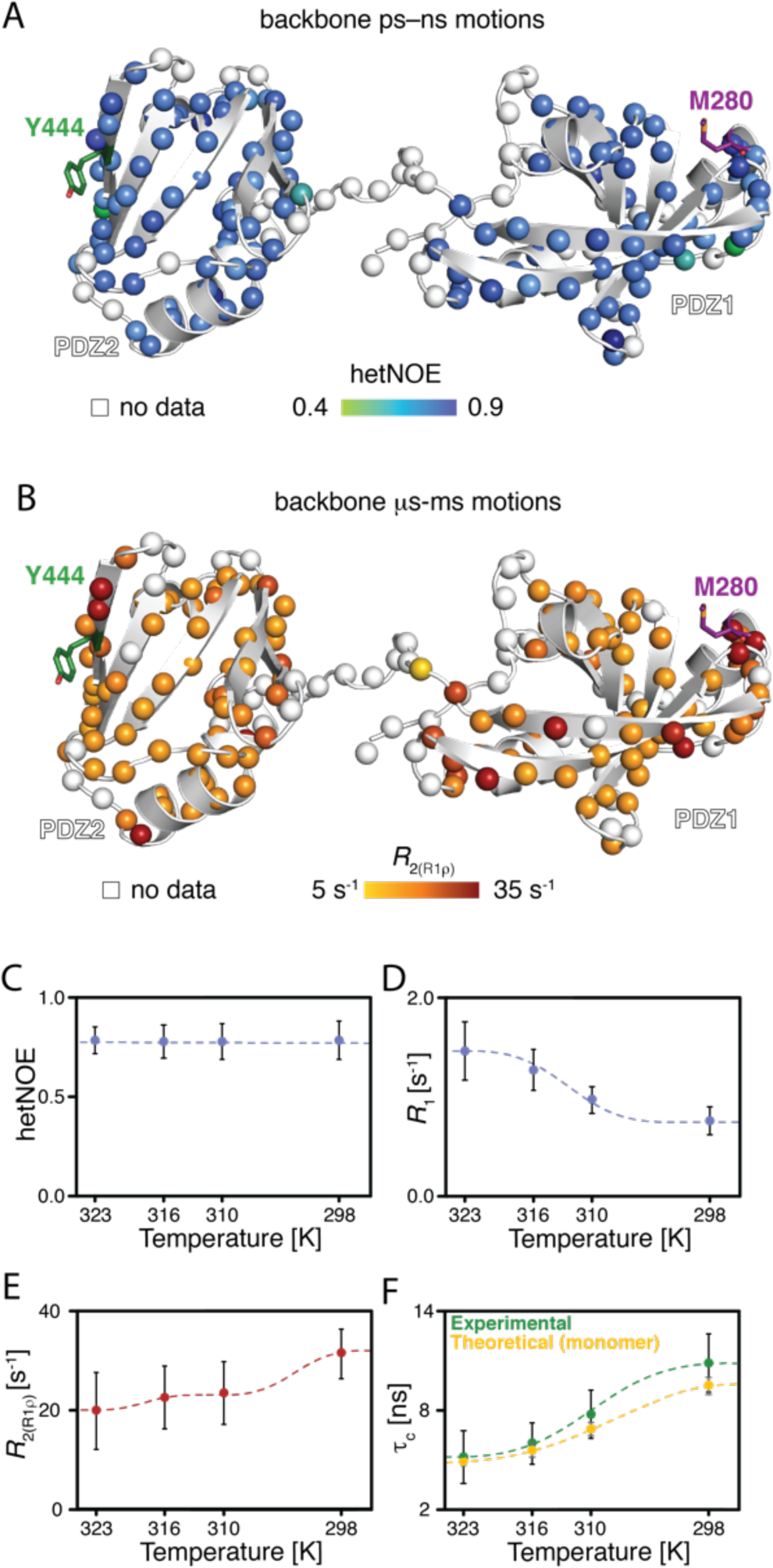
Backbone dynamics of the PDZ1–PDZ2 construct. (**A**) Local backbone dynamics on the ps– ns timescale probed by hetNOE measurements at 50°C. The amide moieties of the PDZ1–PDZ2 construct are shown as spheres and the hetNOE values are indicated by the green to blue gradient. (**B**) Obtained transversal relaxation (*R*_2(R1ρ)_) rates reporting on microsecond motions at 50°C, are plotted on the amide moieties and are indicated by the yellow to red gradient. (**C–E**) Temperature dependence of the average hetNOE (**C**) *R*_1_ (**D**) and *R*_2_-rates (**E**) over the indicated temperature range. Error bars are the S.D. (**D**) Temperature-dependence of the measured rotational correlation time, *τ*_c_, over the indicated temperature range (green). Theoretical values were calculated for the DegP crystal structure (PDB-ID: 3OU0) with HYDRONMR (*68*) for a PDZ1–PDZ2 monomer.

Subsequently, we analyzed the contributions of motions in the micro- to millisecond regime. For these slow timescale motions, we analyzed the ^15^N transverse relaxation rates. First, we measured the *R*_2_-rates derived from the *R*_1_*_ρ_* rates (*R*_2(R1_*_ρ_*_)_), which report on the motions on the lower micro-second timescale, because under the used spin-lock radio frequency (RF) field of 2,000 Hz all exchange contributions (*R*_ex_) slower than 80 µs would be leveled out (Fig. 4B and fig. S8B). In line with the previous analysis of the fast timescale motions, we observed a largely planar profile for the folded segments. In addition, we observed slightly enhanced transversal relaxation rates for the loop regions and for helix *α*_1_ as well as the carboxy-terminal strand *β*_10_, containing M280 and Y444, respectively. Furthermore, we measured the transverse relaxation of the slowly relaxing ^15^N[^1^H] doublet component (*R*_2_*_β_*), revealing exchange contributions (*R*_ex_) on the higher micro- to millisecond timescale (*33*), which resulted in similar profiles as observed for *R*_2(R1_*_ρ_*_)_. In summary, the transversal relaxation rates indicate the existence of local conformational exchange contributions in the regions involved in the interdomain lock.

Next, we assessed the obtained relaxation rates globally in a temperature dependent manner (Fig. 4C–F). Whereas we observed no differences in the hetNOE values over the tested temperature range, we could observe temperature dependent modulations for *R*_1_ and *R*_2(R1_*_ρ_*_)_ resulting consequently in changes in the rotational correlation time *τ*_c_(Fig. 4F). These observed changes in *τ*_c_ can be attributed to the following factors: changes in temperature and consequently solvent viscosity *via* the Stokes Einstein equation lead to a change by a factor of 1.74, eventual decoupling of the individual domains, and/or transient intermolecular interactions. To delineate if other contributions besides the temperature dependent modulation play a role for the observed changes in *τ*_c_, we calculated the theoretical diffusion coefficients and compared them to our experimental data (Fig. 4D). There was a very good agreement at the higher temperatures, ruling out a meaningful contribution by domain decoupling under these conditions, whereas at 25°C the experimental and theoretical values deviate (the experimental values indicate a larger molecule size), pointing towards the formation of a transient interdomain interaction. This observation is therefore in complete agreement with a temperature dependent transition as initially observed within the hexamer-trimer transition for full-length DegP^S210A^ by NMR diffusion measurements (Fig. 1D–F). To obtain a more detailed picture of the roles of individual amino acids involved in the PDZ1–PDZ2 interaction we employed the Lipari-Szabo model-free approach (*34*). This analysis using an axially symmetric diffusion model *D*_||/⊥_ ranging from 1.25 – 1.35 (table S2) over the used temperature range revealed distinct temperature-dependent differences for the two domains (Fig. 4A) for the nanosecond timescale order parameters S^2^, reporting on the fast motions of the N–H vector. Whereas the core of PDZ1 (e.g. central *β*-sheet) was showing only slightly temperature modulated S^2^ values of ∼0.7, the core of PDZ2 showed lower values, with a minimum achieved at 43°C with ∼0.6 for its central *β*-sheet (Fig. 5A and fig. S8C). The exchange rate contributions, *R*_ex_, additionally determined by the Lipari Szabo approach (*34*), are indicative for possible effects of conformational exchange on the micro– to millisecond timescale. In agreement with the initially determined transversal relaxation rates, we observed enhanced *R*_ex_-rates for the residues residing in helix *α*_1_ and parts of the *β*-sheet core of PDZ1 pointing towards dynamical adaptions enabling client interaction (Fig. 5B and fig. 8C). Assessing the *R*_ex_-rates in a temperature-dependent manner reveals that at 25°C possible conformational exchange contributions are minimized, especially around M280, whereas from 37°C onwards micro- to millisecond motions manifest (Fig. 5B, C). Based on the observed behavior the picture emerges, that the extend of motions of M280 is coupled to the release of the inter domain lock and in this way directly modulates the degree of motions within the whole PDZ1 domain (Fig. 5D and fig. S8 C, D). Nevertheless, the observation that domain opening and enhanced backbone dynamics occur on different timescales indicates distinct separate underlying processes, such as domain release and substrate binding.

**Fig. 5:**
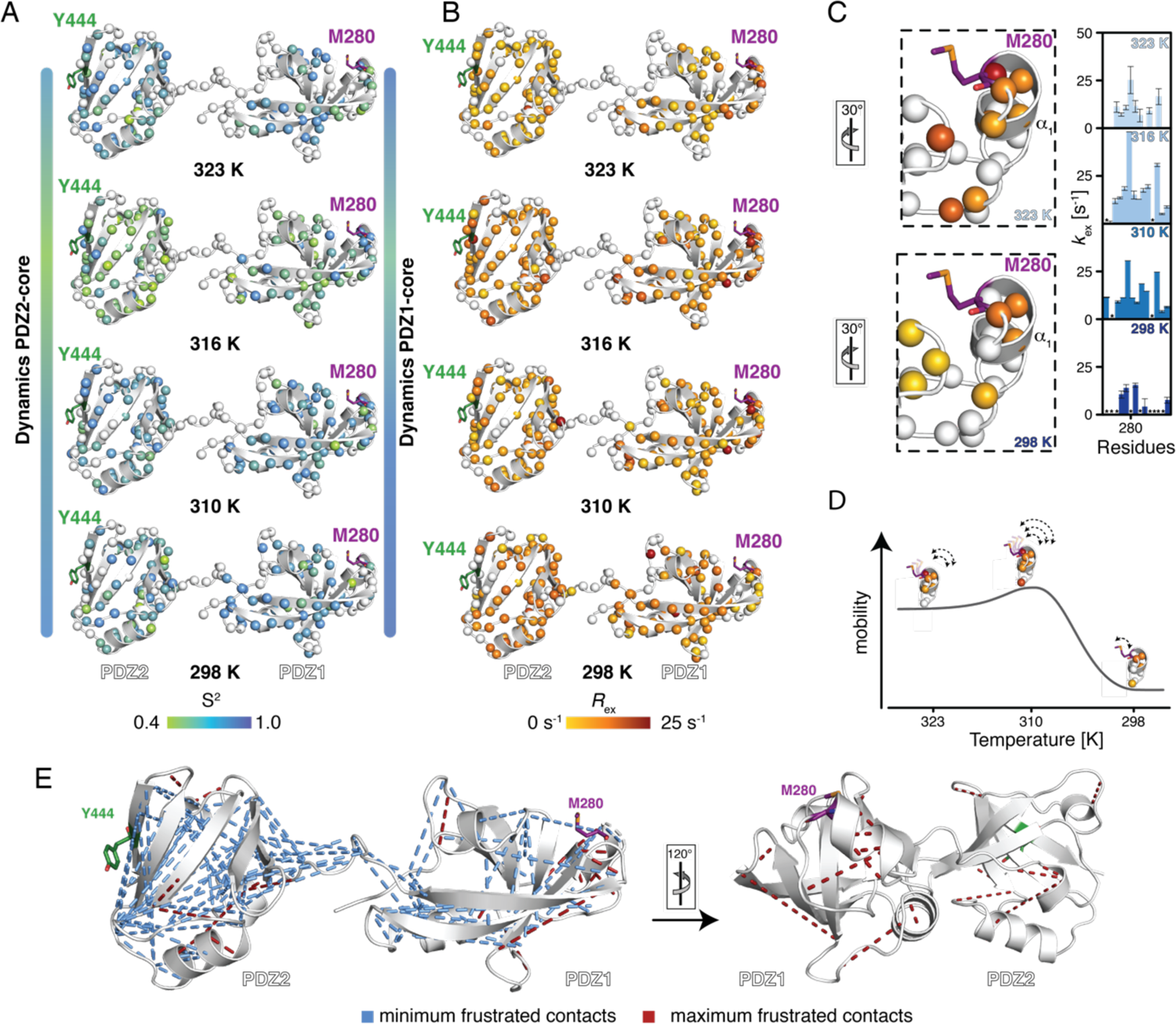
Backbone dynamics show correlated movements of PDZ1 and helix *α*_1_ harbouring M280. (**A, B**) Temperature dependence of the sub-nanosecond order parameter S^2^ (**A**) and the exchange rate *R*_ex_ (**B**), reporting on the flexibility on the micro- to millisecond timescale, calculated using the Lipari-Szabo model-free approach (*34, 36*). (**C**) Temperature dependence of the *R*_ex_-rates and zoom-in on the region around Met280. The rotation relative to panel **B** is indicated. The excerpt from the obtained exchange rates in this region is also shown. Residues for which no data could be obtained are indicated with a *. The full data set is provided in fig. S8C. (**D**) Cartoon representation of the temperature-dependence of the inherent mobility as reflected by the possible conformational exchange contributions on the micro to millisecond timescale of helix *α*_1_ containing M280. (**E**) Frustration analysis was calculated for the DegP crystal structure (PDB-ID: 3OU0) *via* the Protein Frustratometer 2 (*37*). Minimally frustrated interactions are indicated as blue lines, whereas highly frustrated interactions as red lines.

As the domain release apparently had a direct effect on the dynamics of the PDZ1, we wondered what might be the driving force for locking the domains in the inactive form. As for example, molecular chaperones were previously shown to recognize locally frustrated regions (*35, 36*), we hypothesized that local frustration within parts of the PDZ1–PDZ2 domains (*37*) might be the driving force for the observed intermolecular PDZ1–PDZ2 interaction (Fig. 5E). Indeed, the most frustrated region as evidenced by the large number of highly frustrated contacts within the PDZ1–PDZ2 construct resides in helix *α*_1_ harboring M280. As local frustration arises from conflicts between the constrained connectivity of amino acids, like hydrophobic amino acids in close proximity to highly charged residues, resulting in energetically unfavorable local regions, this observation indicates that the M280–Y444 lock reduces this energetically unfavorable state in cellular ground states. In line with this reasoning, both the sub-nanosecond order parameter S^2^ as well as the exchange contributions *R*_ex_ determined *via* the Lipari-Szabo approach indicate the least extent of motions at the lower temperatures. In contrast under heat shock conditions at 43°C the largest degree of inherent motions is observed, pointing to a preparation of the PDZ1 for substrate interaction by modulating these dynamic properties in a temperature-dependent manner.

### Dynamics of the methyl side-chains

To derive the origin of the enhanced local exchange contributions of helix *α*_1_, that provide the basis for modulating the PDZ1–PDZ2 interaction, we exploited the increased sensitivity of methyl groups to gain insight into side-chain dynamics. We chose specific labeling of isoleucine, leucine, and valine methyl groups as these amino acids are well-dispersed among the PDZ-domains providing specific probes. First, we measured [^1^H,^13^C]-NMR spectra at different temperatures between 25°C and 50°C (fig. S9A, D). Comparing the spectra at 50°C and 25°C revealed that a subset of the resonances underwent severe line-broadening (e.g. L272*δ*1, L272*δ*2, L276*δ*1, L272*δ*2, I304*δ*1, I310*δ*1, I318*δ*1, I371*δ*1, I408*δ*1, V431*γ*1, V432*γ*2) accompanied by chemical shift perturbations (fig. S9A–F). Whereas the latter can be attributed to a slight change in compactness of the protein core, which is also evidenced by the decreased extent of global backbone dynamics at the lower temperature (Fig. 5D), the lack of signal intensity for a subset of methyl groups indicates the presence of an intermolecular interaction. To assess the nature of the underlying motions we first determined the product of the side-chain order parameters and the correlation time of the overall molecular tumbling (S^2^•*τ*_C_) for an isoleucine-labeled as well as a leucine, valine-labeled PDZ1–PDZ2 sample, providing insight into the extent of the amplitude of motions of the methyl axis in the picosecond regime (Fig. 6A, B). The obtained residue-wise profiles at 37°C indicate a large degree of flexibility for the studied methyl-groups. Whereas groups residing in the core of the PDZ2-domain show overall a lesser extend of motions on this timescale, reflected by an average S^2^•t_C_ = 4.2 ± 1.6 ns, methyl groups in the core of the PDZ1 uniformly show a higher degree of flexibility with S^2^•t_C_ = 3.0 ± 1.3 ns.

**Fig. 6:**
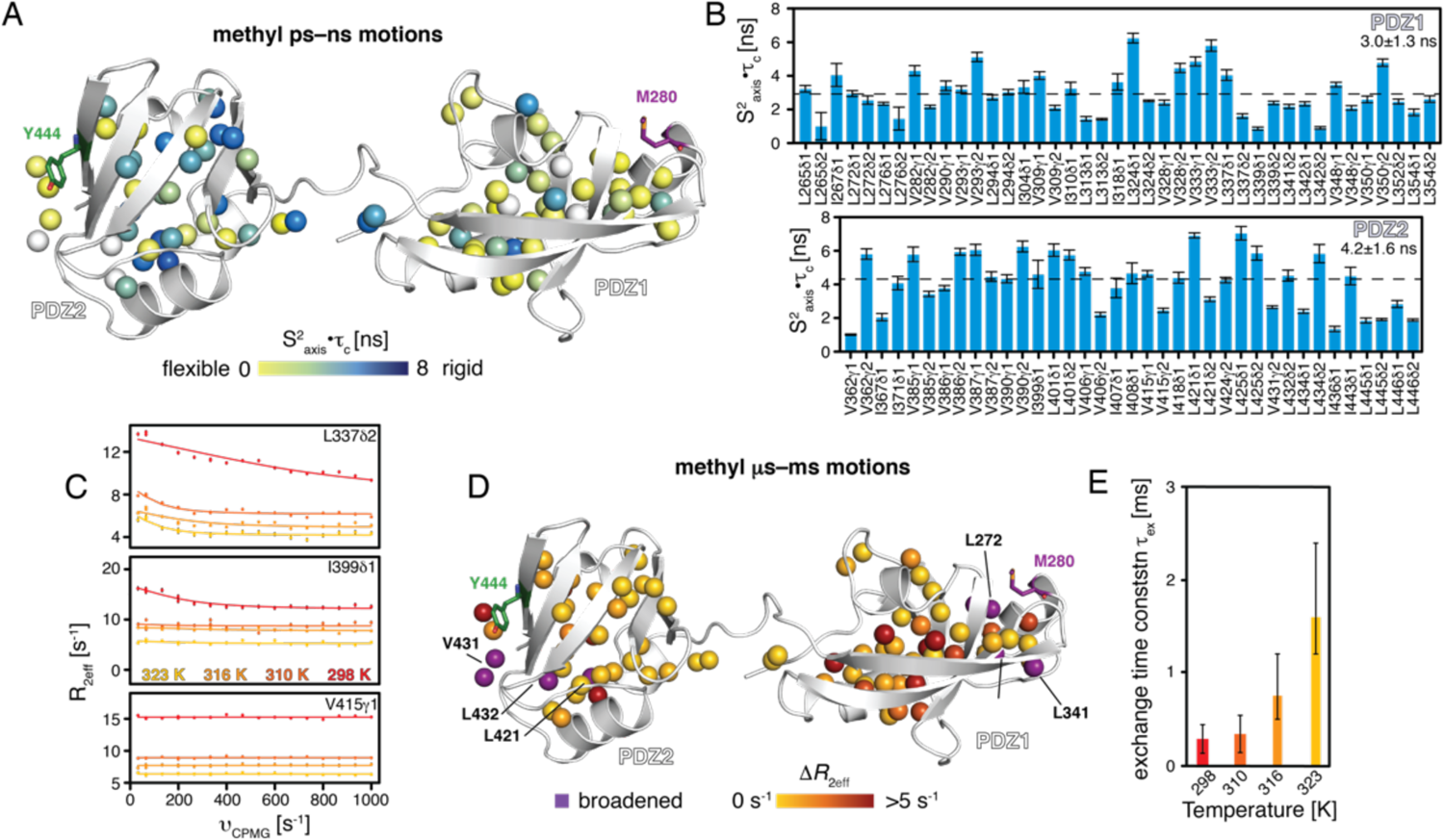
ILV methyl groups reveal conformational exchange within the PDZ1 core. (**A, B**) Local methyl-group dynamics on the pico- to nanosecond timescale probed by methyl triple-quantum (TQ) relaxation experiments measurements (*69, 70*) at 37°C showing the product of the local order parameter and the overall tumbling constant, S^2^_axis_•*τ*_C_. The ILV methyl groups of PDZ1–PDZ2 construct are shown as spheres and the obtained S^2^_axis_•*τ*_C_-values by the green to blue gradient (**A**). S^2^_axis_•*τ*_C_ values plotted against the sequence (**B**). (**C**) Exemplary ^13^C methyl MQ-CPMG relaxation dispersion profiles at different temperatures as indicated. Non-flat profiles indicate millisecond-dynamics. (**D**) Structural view of the amplitude of the CPMG relaxation dispersion profiles *ΔR*_2eff_ at 25°C. (**E**) Rate constants of the dynamic process (*τ*_ex_) obtained from a global fit of the CPMG relaxation dispersion data.

As several methyl resonances appeared to be involved in conformational exchange processes, we next quantified these by employing multiple quantum Carr-Purcell-Meiboom-Gill (CPMG) relaxation dispersion measurements (*38*), sensing motions on the micro– to millisecond timescale. Measuring CPMG experiments at temperatures ranging from 25–50°C revealed that the residues experiencing micro- to millisecond motions reside mainly within the PDZ1 domain, namely methyl-groups V282*γ*2, V309*γ*2, I310*δ*1, L313*δ*1, I318*δ*1, L337*δ*2, L339*δ*2, L341*δ*1, L352*δ*2, L354*δ*2, as evidenced by non-flat dispersion profiles (Fig. 6C–E and fig. S9G). Quantitative global analysis of the dispersion data showed that this motion occurred on a 0.2 ms (25°C) or 0.4 ms (37°C) timescale under inactive conditions and 1 ms (42°C) or 2 ms (50°C) timescale under activating heat shock conditions (Fig. 6E). The observation of the increased timescale of the side-chain motions under increased temperatures indicates pronounced modulations of inherent side-chain dynamics, which is directly coupled to the release of helix *α*_1_, suggesting contributions of the local frustration driving this dynamical behavior (Fig. 5E).

**Fig. 7:**
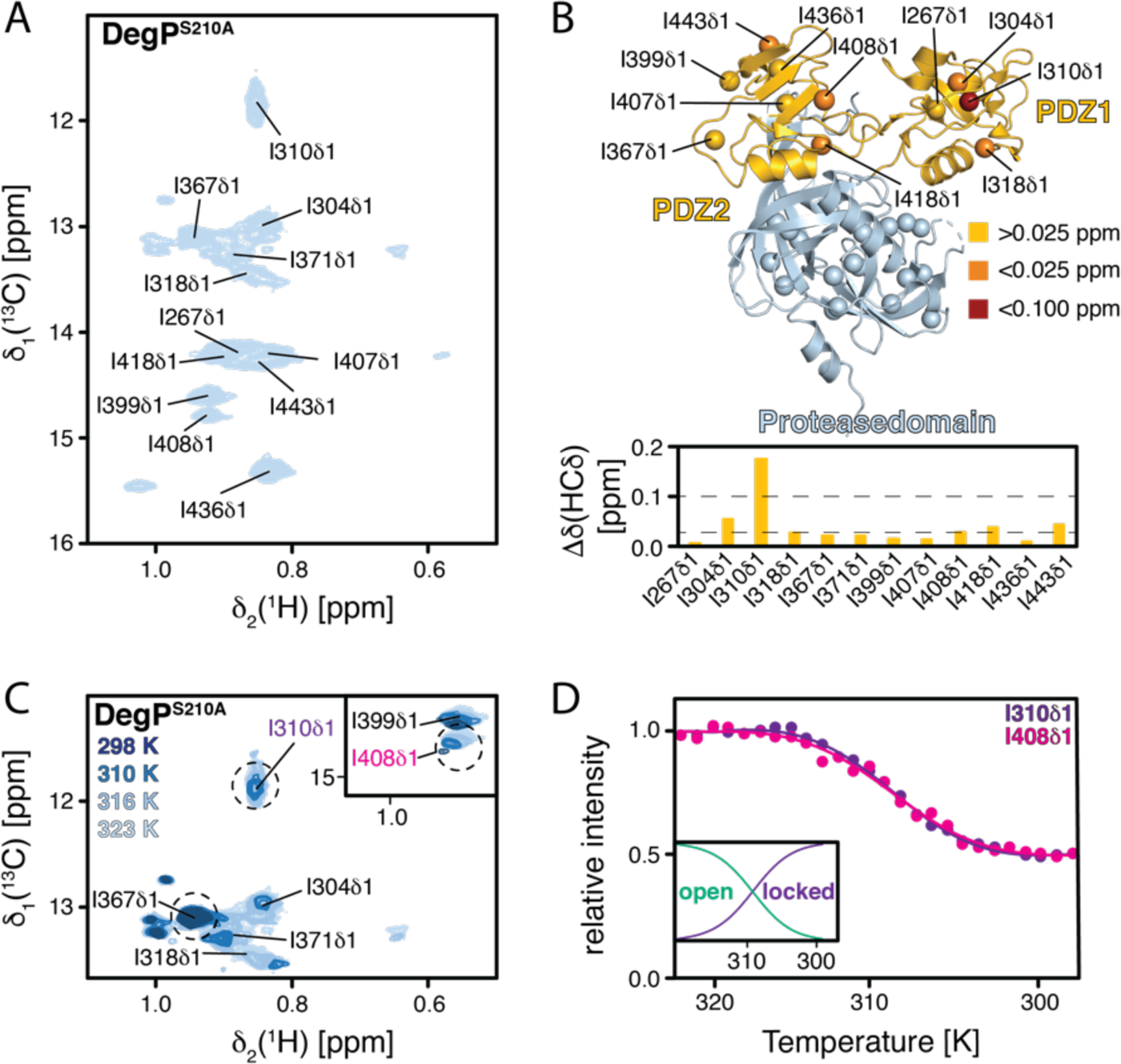
Isoleucine methyl groups within full-length DegP show also temperature-dependent transition. (**A**) 2D [^13^C,^1^H]-NMR spectrum of [*U*-^2^H, Ile-*δ*_1_-^13^CH_3_]–DegP^S210A^ at 50°C. The indicated resonance assignments were transferred from the assigned PDZ1–PDZ2 construct. (**B**) Chemical shift changes of the isoleucine methyl groups of DegP compared to the PDZ1–PDZ2 (bottom). The isoleucine methyl groups of one DegP monomer are indicated with spheres (top), and the chemical shift changes compared to the PDZ1–PDZ2 construct are indicated by the yellow to red gradient. (**C**) Excerpt of the isoleucine methyl region of a 2D [^13^C,^1^H]-NMR spectrum of [*U*-^2^H, Ile-*δ*_1_-^13^CH_3_]–DegP^S210A^ at different temperatures as indicated. (**D**) The temperature dependence of the selected I310*δ*1 and I408*δ*1 signal intensities is shown. Solid lines represent a fit to a 2-state model for the extraction of thermodynamic parameters. The inset shows the respective populations of the open and the locked state, respectively, at the different temperatures.

Having established the inherent flexibility on the micro- to millisecond timescale of mainly the PDZ1 domain likely crucial for substrate recognition, we next set out to probe these effects on the isoleucine methyl probes within the full-length DegP^S210A^. Based on the side-chain assignment of the isolated PDZ1–PDZ2-domain construct (fig. S2E), we were able to unambiguously transfer the assignment of all eleven resonances within the DegP^S210A^ full-length construct (Fig. 6A, B). In order to study the transformation from the low temperature hexameric-state to the high-temperature trimeric-state within full-length DegP^S210A^, we measured [^1^H,^13^C]-NMR spectra over the complete temperature range from 25°C to 50°C (Fig. 6C). At the low temperatures, several residues (e.g. I310*δ*1, I408*δ*1) show reduced intensity comparable to the situation observed for the PDZ1–PDZ2 construct. Using the measured intensities for I310*δ*1 and I408*δ*1 from 25°C to 50°C together with transition state theory, we extracted thermodynamic values for the transition from the locked to the open state. Fitting the data to a 2-state model yielded an activation enthalpy of *Δ*H=241.7 ± 88 kJ mol^-1^ and entropy of *Δ*S=0.78 ± 0.14 kJ mol^-1^ K^-1^ with a transition midpoint at 36.8 ± 1.3°C. The obtained activation enthalpy is perfectly in line with the obtained values for a general methyl–phenylalanine or a comparable methionine–phenylalanine interaction motifs of 50–100 kJ mol^-1^ (*39, 40*), providing further indication of the Met–Tyr motif stabilizing the intermolecular PDZ1–PDZ2 interaction.

Similar to the initial observations for the [^1^H,^15^N]-NMR spectra (fig. S1), the [^1^H,^13^C]-NMR spectra of the isoleucine labeled DegP^S210A^ also did not yield high-quality spectra for the protease domain of full-length DegP (Fig. 6). This suggests that a high-amount of inherent dynamics exists for this domain (backbone as well as methyl groups), which could already be observed to cause poor quality NMR spectra for some oligomeric proteases earlier (*41, 42*), and therefore likely reflects the regulatory role of the variety of different loops involved in the regulation of DegP protease function (fig. S1A, B). In agreement with this reasoning is the observation of spectra of similar quality for the [^1^H,^13^C]-NMR spectra of isoleucine labeled DegP^S210A,Y444A^ and DegP^S210A,M280I^ over the tested temperature range (fig. S10A–C). Whereas these spectra differ in terms of quality for the resonances of the two PDZ domains, with DegP^S210A,Y444A^ providing the highest quality spectra, reflecting its obligate trimeric form over the whole temperature range assessed, no distinctive differences for isoleucine residues originating from the protease domain could be observed.

### DegP trimers display increased proteolytic activity

To investigate if DegP trimerization has a direct influence on the proteolytic function, we tested the effect on previously reported DegP model substrates: fluorescent p23 peptide as well as β-casein. In initial test experiments, we were not able to observe any significant differences for the DegP variants cleaving the fluorescent p23 peptide (fig. S11), which can likely be attributed to the proposed function of this peptide as a DegP activator (*23*). On the other hand, the used DegP mutants, DegP^Y444A^, DegP^M280A^, and DegP^M280I^ did cleave β-casein more rapidly compared to DegP in a coupled assay using a non-activating fluorescent reporter peptide (*14*) (Fig. 8A). The observation of sigmoidal curves for all tested DegP variants indicates the necessity of a second activation step, besides breakage of the PDZ1–PDZ2 interaction, for its proteolytic function, likely pointing to the positive cooperative allosteric activation of the LA-Loop upon β-casein binding to PDZ1 (*12, 18, 23*). Interestingly, the DegP^M280I^ variant showed an intermediate behavior in our essays, suggesting a partially functioning interdomain lock, albeit being surprisingly active at 25°C compared to 37°C.

**Fig. 8:**
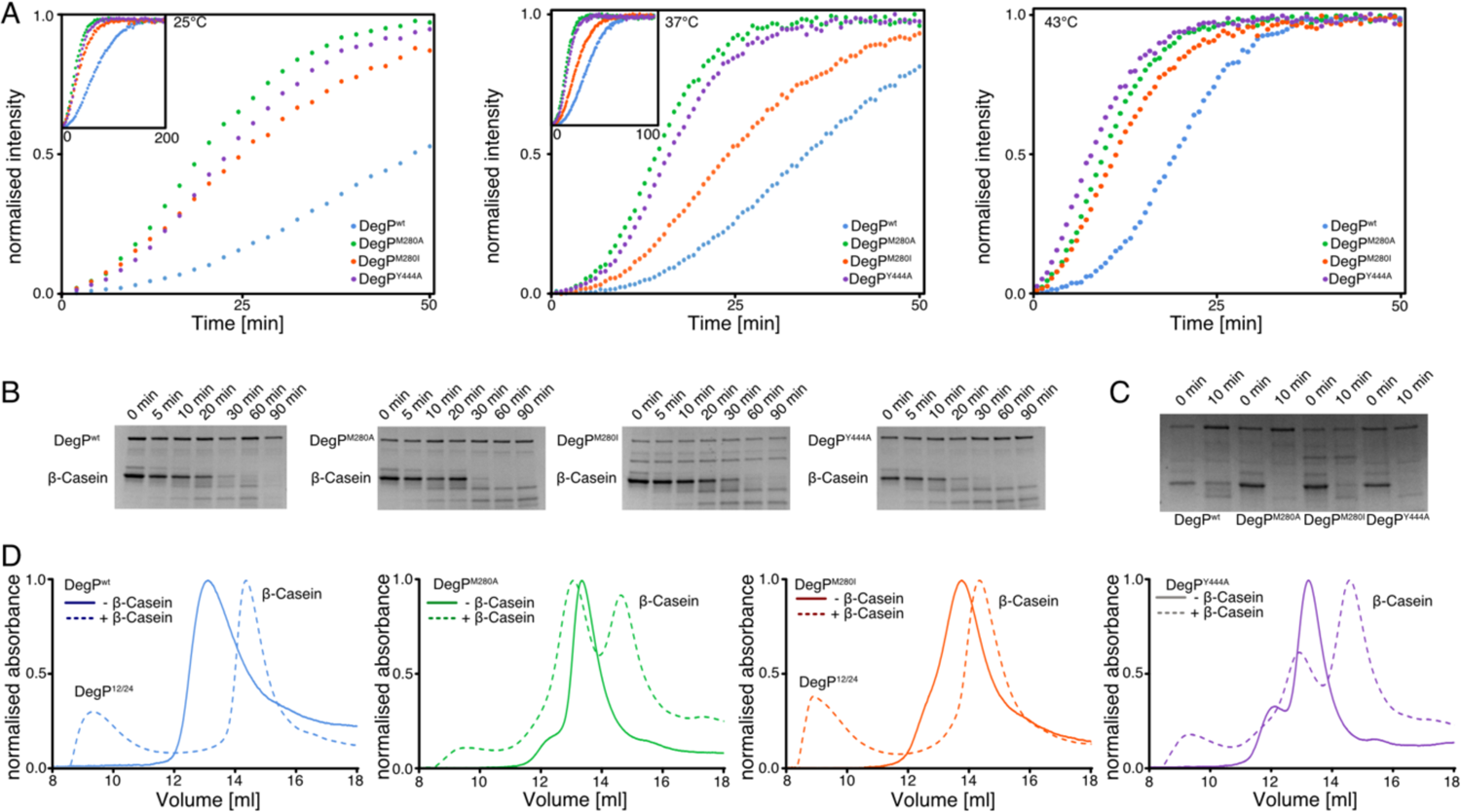
Assessing the effect of the domain-lock on DegP proteolytic function. (**A**) DegP (10 µM monomer concentration) proteolysis of β-casein (50 µM) *via* detection by a fluorescent reporter peptide (100 µM) at 25°C, 37°C, and 43°C. All experiments were scaled to the experiment at 43°C. Insets show the respective full data sets at the lower temperatures. Experiments were performed as triplicates yielding identical results. (**B**) SDS-PAGE analysis of β-casein (5 µM) cleavage by DegP variants (1 µM monomer concentration) at the indicated time intervals at 25°C. (**C**) SDS-PAGE analysis of β-casein cleavage at the indicated time intervals at 37°C. Experiments were performed as triplicates yielding identical results. (**D**) Size-exclusion chromatography elution profiles of DegP and β-casein mixtures on an S200increase 10/300 column (GE Healthcare).

Analyzed by SDS-PAGE, DegP^M280A^ and DegP^Y444A^ variants completed cleavage of a 30-times molar excess (considering a DegP hexamer) of β-casein within 30 min, whereas DegP and DegP^M280I^ required up to 1 hour to process β-casein (Fig. 8B, C). It should be noted that it should be possible for DegP^M280I^ to form 12- and 24-meric cages due to interactions between the long methyl-bearing isoleucine side-chains and the aromatic π-system of Y444 (*29*). This feature could be confirmed by size-exclusion chromatography, where DegP and DegP^M280I^ formed almost exclusively cage-like structures, reflecting DegP cages in the presence of the β-casein substrate, whereas the capability to assemble these cages was largely diminished in DegP^M280A^ and DegP^Y444A^ indicating the importance of the PDZ1-PDZ2 interdomain stabilizing the large assemblies (Fig. 8D).

Additional NMR titrations provided evidence for our initial reasoning, that p23 primarily binds to the PDZ1 domain and thus likely triggers the allosteric changes needed for proteolytic activation *via* the LA-loop in complete agreement with previous crystallographic studies (*23*) (fig S10A–D). β-casein on the other hand had a more global effect on both domains at half equimolar ratio, reflecting its larger size (24 kDa) and consequently larger binding interface compared to the p23 peptide (fig. S11D). At lower ratios, the largest chemical shift perturbations were observed for the PDZ1 domain, in agreement with its role as the main substrate recognition region and indicating an auxiliary role for the PDZ2 in substrate recognition (fig. S11D, E). Finally, the non-activating reporter peptide did not lead to any significant spectral changes, indicative of binding to the protease domain (*43*) and thus highlighting the need for allosteric activation of DegP through substrate binding to the PDZ1 domain, a characteristic feature of a variety of HtrA proteins (*11, 14, 20, 44, 45*).

## Discussion

Despite being studied extensively and the existence of several high-resolution crystal structures of DegP in its different states multiple questions remain about how its structural transitions are achieved. Within the known structures of the hexameric state, the DegP_12_ and DegP_24_ structures, an intermolecular PDZ1–PDZ2 interaction always coordinates the arrangements of the DegP trimer building-blocks to stabilize the larger oligomeric assemblies (*12*). Thereby, the structures indicated a different arrangement of the PDZ-domains for the hexameric state compared to the larger oligomeric states (Fig. 1A). However, our solution NMR results presented here shows that in solution DegP is locked in the hexameric state *via* the same M280– Y444 interaction as observed initially within the larger oligomeric states, indicating that this interaction is crucial for modulating the different oligomeric states within DegPs functional cycle. Although the participation of Y444 was already suggested based on the characterized DegP^S210A,Y444A^ mutant (*14*), our data revealed that the M280 is not only participating in this interaction but effectively modulating the temperature-dependent opening of this interdomain lock.

Studying backbone in combination with methyl-group side-chain dynamics over the transition temperature range revealed inherent dynamics within the core of the PDZ1 domain (V282, V309, I310, L313, I318, L337, L339, L341, L352, L354), which are likely allosterically triggered by the domain release. The conformational transition of M280 between its locked and open state is on a different timescale therefore only indirectly modulating these inherent faster dynamics. Our data indicate that this interaction is stabilized likely *via* a sulfur-*π* interaction of the M280 and Y444 side-chains, a motif lately receiving recognition as an important stabilizer of protein interactions (*28, 39*). Although we cannot completely rule out a methyl-π interaction, the fact that a sulfur-π is believed to be the stronger interaction (*39, 40*) and our observation that the conservative mutation M280I leads to a destabilization of the inter-domain lock suggests a sulfur-π interaction.

Taken together our backbone and side-chain dynamical data draw a detailed picture of how the temperature-dependent breakage of the interdomain lock fine-tunes the inherent dynamics of PDZ1 to ensure efficient substrate interaction (Fig. 9). Whereas the lack of backbone motions at 25°C directly explains the consequence of the locked conformation at low temperatures, likely driven by a reduction of the local frustration in M280 bearing helix *α*_1_ in cellular ground states, the situation changes with increasing temperatures, where we begin to observe extensive inherent dynamics. The amplitude of motions at 37°C (Figs. 5A and 6D) reveals a fine-tuned dynamical compensation, which can easily adapt to slight changes in temperature to either lock the interaction (at a lower temperature) or unlock the interaction and consequently activate also the protease function of DegP (at a higher temperature). Whereas under heat-shock conditions the extensive inherent dynamics on the micro- to millisecond timescale within PDZ1 indicate a domain optimized for substrate interaction facilitating subsequent proteolytic cleavage by the protease domain. This finding poses the very interesting question whether and how the proposed large DegP-cages can form at elevated temperatures. Therefore, under heat-shock conditions the formation of the larger DegP states might not be supported by the stabilization of the PDZ1–PDZ2 lock, even though cages are supposed to form between 25–48°C (*23*). Alternatively, upon cage-formation there could be an allosteric feedback loop from the protease domain back towards the PDZ1 domain, with the resulting compensation of the frustrated area around *α*_1_ once again forming the main driving force for a stable complex formation *via* the M280–Y444 domain-lock (Fig. 9). The exact nature of the existence and/or the stabilization of the interdomain lock at elevated temperatures clearly needs further research as most studies so far focused on lower temperatures (*12, 15, 17, 23*).

**Fig. 9:**
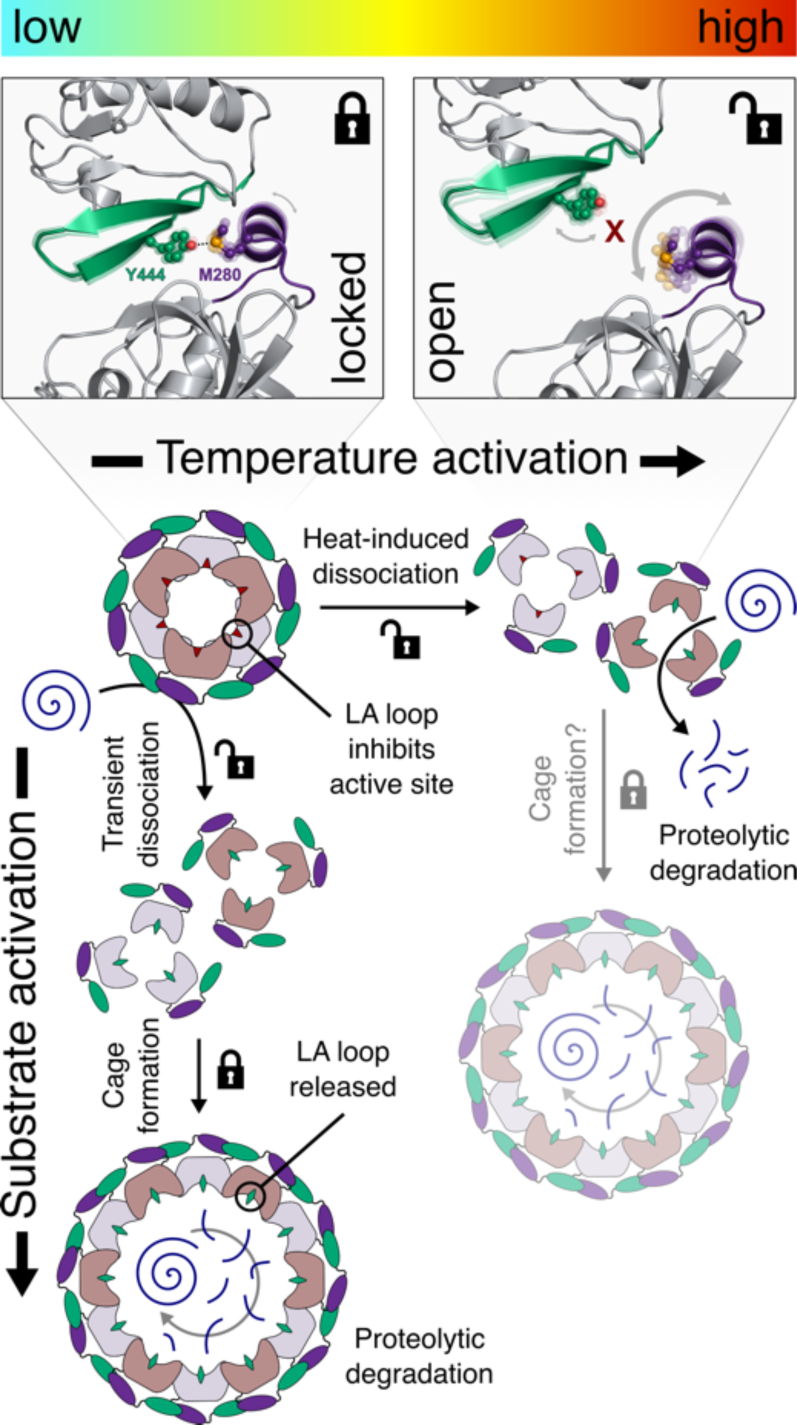
Temperature activation of the DegP protease mediated by the M280–Y444 interdomain lock. In cellular ground-states at low temperature the dynamics of M280 are reduced to stabilize the PDZ1–PDZ2 interaction *via* the sulfur-*π* interaction with Y444 (locked state), resulting in an inactive DegP hexamer. Under heat shock conditions, M280 exhibits a large degree of flexibility and the interaction with Y444 is broken (open state), as it cannot compensate the overall increased dynamics and the PDZ1–PDZ2, leading to trimerization of DegP. At low temperatures the protease activity of DegP can be triggered by activating substrates, which mediate the formation of proteolytic (12/24-meric) cages *via* a transient trimerization through interactions with the PDZ1 domain. Upon temperature activation the M280–Y444 lock mediates the transformation of DegP from the locked inactive hexameric state to an open trimeric state, primed for the proteolytic degradation of substrates, but still requiring activation through substrate binding. Under heat-shock conditions, the structural basis for cate formation remains unclear, even though DegP is still able to form cages under these conditions (*23*).

On a more broader scope, the involved carboxy-terminal Y444 remarkably is found in several HtrA proteins known to exploit a similar locking mechanism (fig. S12A–E) (*12, 21, 46, 47*), Furthermore, the involved residues in DegP, M280 and Y444, are also highly conserved within different HtrA-proteins carrying two PDZ-domains, with the tyrosine showing a slightly higher conversation score than the methionine (fig. S12F, G). Our finding might therefore be indicative of a more wide-spread mechanism not only restricted to HtrA-proteins as it is already known that carboxy-terminal aromatic/hydrophobic residues in a variety of mammalian multi-PDZ domain proteins govern their formation of homo- and heterodimers suggesting a possible evolutionary conserved mechanism of stabilizing PDZ–PDZ interactions (*48, 49*).

## Materials and Methods

### Cloning of different DegP domain constructs

The pDS4 plasmid contains wild-type DegP (*E. coli* K12 strain) lacking its amino-terminally signaling sequence within a pET28b-vector backbone yielding DegP with an amino-terminal His6-Tag (purchased from GenScript). The individual PDZ domains, the PDZ1–PDZ2 construct as well as the SUMO-tag were amplified by PCR from pDS4 and pET28_SUMO_Hsc70 (a kind gift of B. Bukau (Heidelberg)) (*50*) plasmids, respectively. The different DegP-PDZ constructs and the SUMO-tag were fused by standard PCR methods and cloned into a pET28a(-) vector *via* the NcoI and XhoI restriction sites by standard cloning techniques yielding PDZ constructs fused to an amino-terminal His6-SUMO-tag. Plasmids and primers used in this study can be found in table S3.

### Protein expression and Purification of PDZ1–PDZ2 domains

pDS10 (DegP^261-354^), pDS11 (DegP^359-448^), and pDS12 (DegP^261-448^) plasmids were chemically transformed into *E. coli* BL21 Star^TM^ (*λ* DE3) cells. The transformed cells were grown at 37°C in 1 l LB medium containing 50 µg/ml kanamycin until an OD_600_ ≈ 0.5 was reached. Expression was induced by the addition of 0.4 mM IPTG and the cells were left to grow overnight at 25°C.

Cells were harvested by centrifugation at 8,000*g for 20 min at 4°C and subsequently resuspended in 25 ml of lysis buffer (25 mM HEPES/NaOH, 0.5 M NaCl, 10 mM Imidazole, pH 7.5). The resuspended cells were incubated on ice with one Complete EDTA-free protease inhibitor (Roche) tablet as well as with 100 U of DNAse (ArcticZymes) for 30 min and subsequently lysed with a Q700 sonicator (Qsonica) (10 s on, 30 s off, 30 % power). Cell debris was removed by centrifugation at 18000*g for 45 min at 4 °C and the supernatant was applied to a Ni^2+^–Sepharose 6 Fast Flow (GE Healthcare) loaded gravity column, followed by step-wise elution with 20 ml of lysis buffer supplemented with 100 mM and 500 mM imidazole, respectively. Fractions containing the target proteins were dialyzed against PBS (pH 7.4) and the His_6_-SUMO-tag was removed by enzymatic cleavage using human SENP1 protease at 4°C overnight (Addgene #16356) (*51*). The cleaved proteins were applied again to a Ni^2+^–column and the flow-through was collected. The proteins were concentrated using Amicon centrifugal filters (5k MWCO or 10k MWCO, Millipore) and purified further by size exclusion chromatography (Superdex 75 Increase, GE Healthcare) in NMR buffer (25 mM potassium phosphate, 1 mM EDTA 1 mM TCEP, pH 7.0).

For His_6_-tagged full-length DegP variants, an additional unfolding/refolding step was introduced. To this end, after cell lysis and centrifugation, the supernatant was loaded to a HisTrap HP 5 ml column (GE Healthcare) equilibrated with lysis buffer supplemented with 6 M Guanidinium chloride. For on-column refolding, the bound protein was washed with a 6–0 M Guanidinium chloride linear gradient (15 CV, 0.5 ml/min) overnight and subsequently eluted with a step elution of 100 mM and 500 mM imidazole. In the next step, DegP was purified by size exclusion chromatography (Superdex 200 Increase, GE Healthcare) in NMR buffer.

### Isotope labeling

We obtained [*U*-^15^N] and [*U*-^15^N,^13^C] labeled PDZ-domain constructs by growing the expression cells in M9 minimal media (*52*), supplemented with (^15^NH_4_)Cl and *D*-(^13^C)-glucose, respectively. For full-length DegP, the cells were in addition grown in D2O-based M9 minimal medium, yielding [*U*-^2^H,^15^N] and upon the usage of *D*-(^13^C,^2^H)-glucose [*U*-^2^H,^15^N,^13^C] proteins, respectively. For specific methyl group labeling, we either supplemented D2O-based M9 minimal medium with *D*-(^12^C,^2^H)-glucose and isoleucine precursor (2-Ketobutyric acid-4-^13^C,3,3-d_2_ sodium salt hydrate) 1 hour prior to induction resulting in [*U*-^2^H, Ile-*δ*_1_-^13^CH_3_] or in a similar manner with *D*-(^12^C,^2^H)-glucose and leucine, valine precursor [2-Keto-3-methyl-d_3_-3-d_1_-4-^13^C-butyrate] resulting in non-stereospecific [*U*-^2^H, Leu/Val-^13^CH_3_/^12^CD_2_] labeled DegP-variants. All isotopes were purchased from Sigma Aldrich/Merck.

### NMR spectroscopy

NMR measurements were performed on Bruker Avance III HD 700 MHz or 800 MHz spectrometers, running Topspin 3.5/3.6 and equipped with a cryogenically cooled triple-resonance probe. All experiments were performed in NMR buffer (25 mM potassium-phosphate, 1 mM EDTA, 1 mM TCEP pH 7.0).

For the sequence-specific backbone resonance assignment experiments of DegP^S210A^ constructs and the different PDZ-domain constructs the following experiments were recorded: 2D [^15^N, ^1^H]-TROSY-HSQC (*53*) and for full-length DegP^S210A,Y444A^ the following TROSY-type 3D experiments 3D trHNCA, 3D trHNCACB, and 3D trHNCO (*54*) whereas for the different PDZ-domain constructs standard through-bond 3D HNCA, 3D HNCACB, 3D HNCO, 3D CBCA(CO)NH (*55*) experiments were used. Aliphatic side-chain resonance assignment for the PDZ1–PDZ2-construct was performed based on 2D [^13^C, ^1^H]-HSQC spectra with/without constant time (CT) version, as well as 3D (H)CC(CO)NH, H(CC)(CO)NH, and HCCH-TOCSY-experiments (*55*). In addition, the following NOESY-type experiments with the indicated mixing times were performed: 3D H_all_-H_aro_-C_aro_ with 120 ms mixing time, 3D H_all_-H_ali_-C_ali_ with 120 ms mixing time (*55*), 3D ^13^C_m_-^13^C_m_^1^H_m_ SOFAST NOESY with 50 ms mixing time (*56*). Assignment experiments were performed at 50 °C if not indicated otherwise.

For the quantitative analysis of signal intensities, the amplitudes were corrected by differences in the ^1^H-90° pulse-length, the number of scans, and the dilution factor (*57*). NMR data were processed with a combination of NMRPipe (*58*) and mddNMR2.6 (*59*) as well as analyzed with CARA (*60*).

For titration experiments, 2D [^15^N, ^1^H]-TROSY-HSQC experiments were acquired with 4 scans and 2048 x 256 complex points in the direct and indirect dimensions, respectively. The chemical shift changes of the amide moiety were calculated as follows:

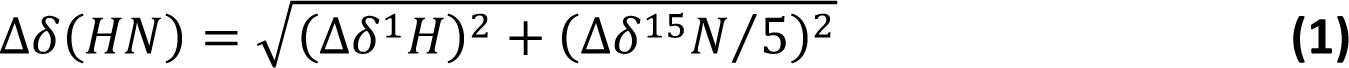

The chemical shift changes of the methyl groups were calculated as follows:

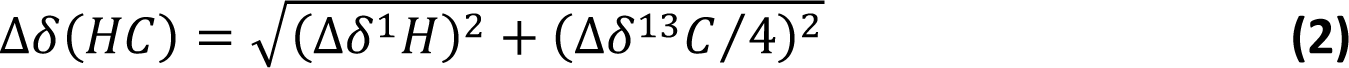

Secondary chemical shifts were calculated relative to the random coil values using the prediction software POTENCI (*61*). Further, a weighting function with weights 1–2–1 for residues (*i-1*)–*i*–(*i+1*) was applied to the raw data (*5, 35*).

### NMR backbone dynamics

For the analysis of the dynamic properties of the different DegP constructs, the following relaxation experiments were measured: ^15^N{^1^H}-NOE (*62*), *T*_1_(^15^N) (*62*), *T*_1_*_ρ_*(^15^N) (*63*), and TROSY for rotational correlation times (TRACT) for determining *T*_2_*_α_*(^15^N) and *T*_2_*_β_*(^15^N) (*24*) at 700 MHz (16.4 T) proton (^1^H) frequency. Non-linear least square fits of relaxation data were done with MATLAB (MathWorks) and the Dynamics Center 2.5 (Bruker Biospin). *R*_2(R1_*_ρ_*_)_(^15^N) values were derived from *T*_1_*_ρ_* (*64*) using equation 3:

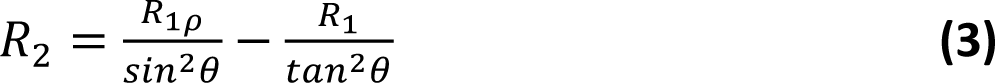

with *Θ = tan^−1^(ω/Ω)*, where *ω* is the spin-lock field strength (2 kHz) and *Ω* is the offset from the ^15^N carrier frequency.

Error bars for *R*_1_(^15^N), *R*_1_*_ρ_*(^15^N), *R*_2α_(^15^N), and *R*_2β_(^15^N) were calculated by a Monte Carlo simulation embedded within Dynamics Center 2.5 (Bruker, Biospin), and for *R*_2(R1_*_ρ_*_)_ (^15^N) by error propagation. Error bars for the ^15^N{^1^H}-NOE were calculated from the spectral noise. Analysis of the obtained relaxation rates was performed with Tensor2 (*65*) using an anisotropic diffusion tensor on the NMRbox web server (*66*). Missing atoms and hydrogens in the PDB structure (3OU0) were added using PDBFixer (*67*). Tmeperature dependent rotation correlation times (*τ*_c_) were estimated by using HYDRONMR version 7c (*68*) *via* NMRBox at 700 MHz (16.4 T) magnetic field strength using a standard AER (atomic element radius) value of 3.3 Å, a 1.02 Å N–H bond length, a CSA of −160ppm, with the refined PDZ1–PDZ2 crystal structure (PDB-ID: 3OU0). For the temperature dependent changes in viscosity of the water, the standard values were obtained from the NIST website (https://webbook.nist.gov/chemistry/fluid/).

### NMR side-chain dynamics

Experiments were performed on either a [*U-*^2^H, Ile-*δ*_1_-^13^CH3]-PDZ1–PDZ2 or a [*U-*^2^H, Leu-*δ*_1,2_-Val-*γ*_1,2_^13^CH_3_]*-*PDZ1–PDZ2 sample at a temperature of 37°C in 99.9% D_2_O based NMR buffer. Side-chain methyl order parameters (S^2^_axis_) were determined by cross-correlated relaxation experiments (*69, 70*). Single-(SQ) and triple-quantum (TQ) ^1^H–^13^C experiments were collected at a series of delay times. Ratios of the peak intensities were fitted for eight values ranging between 2 and 24 ms using the following equation where *T* is the relaxation delay time and *δ* a factor to account for coupling due to relaxation with external protons:

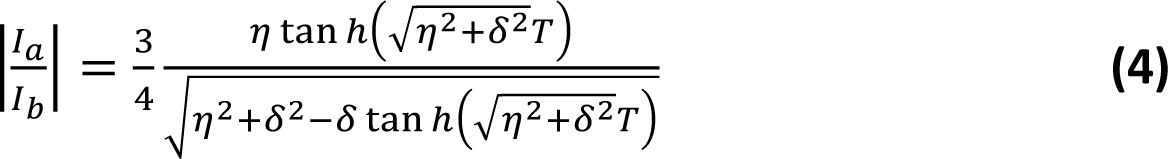

S^2^_axis_ values were determined using equation 5, using their fitted *η* values and the separately determined rotational correlation time adjusted for the change in viscosity in 100% D_2_O (*τ*_c_) of 14.4 ns at 37°C:

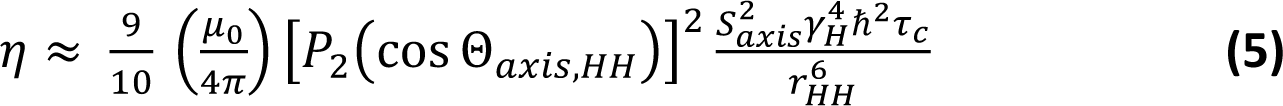

Where µ_0_ is the vacuum permittivity constant, *γ*_H_ the gyromagnetic ratio of the proton spin, *r*_HH_ is the distance between pairs of methyl protons (1.813 Å), S^2^_axis_ is the generalized order parameter describing the amplitude of motion of the methyl 3-fold axis, *Θ*_axis,HH_ is the angle between the methyl symmetry axis and a vector between a pair of methyl protons (90°), and P_2_(x) = ½ (3x^2^-1). Finally, the product of the methyl order parameter and the overall correlation time constant, S^2^_axis_•*τ*_C_, was determined.

Multiple quantum (MQ) methyl relaxation dispersion experiments (*38*) were recorded as a series of 2D data sets using constant time relaxation periods (*T*) of 30 ms (800 MHz (18.8 T); 25°C, 37°C, 43°C, and 50°C) and CPMG (Carr-Purcell-Meiboom-Gill) frequencies ranging from 67 to 1,000 Hz. *R*_2,eff_, the effective transverse relaxation rate was calculated according to the following equation:

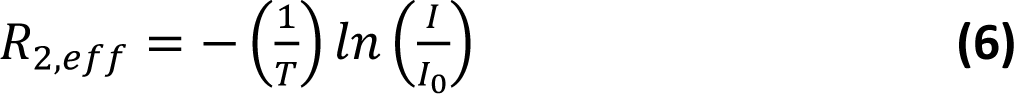

where *I* (or *I*_0_) are the intensities with and without the presence of a constant-time relaxation interval of duration *T*, during which a variable number of ^13^C 180° pulses are applied leading to *ν*_CPMG_ = 1/(2*δ*), where *δ* is the time between successive pulses. Dispersion data were fitted numerically to a two-site exchange model using the program ChemEx (available at https://github.com/gbouvignies/chemex/releases).

### NMR magnetization exchange

2D methyl magnetization exchange experiments were recorded to quantify the exchange between the locked (l) and open (o) state of the Met280-^13^C*ε*^1^H*ε* methyl group as outlined by Farrow *et al.* (*30*). The ^13^C-EXSY-SOFAST-HMQC experiments (*31*) were recorded with varying delays between 50 and 1,000 ms at 37°C. To ensure equilibration of the initial magnetization a long relaxation delay (1s) was chosen. The signal intensities of the observed three peaks of M280-^13^C*_ε_*^1^H*ε*, namely, the auto peak of the locked conformation, *I*_ll_, and the auto peak of the open conformation *I*_oo_ as well as the cross peak originating from the transition from the locked to the open state, *I*_lo_ was determined. Due to signal overlap the second cross-peak, *I*_ol_, could not be determined from the spectra, therefore it was assumed in a first order approximation that it follows a similar build-up than *I*_lo_. All data points were simultaneously fitted to the following theoretical equations:

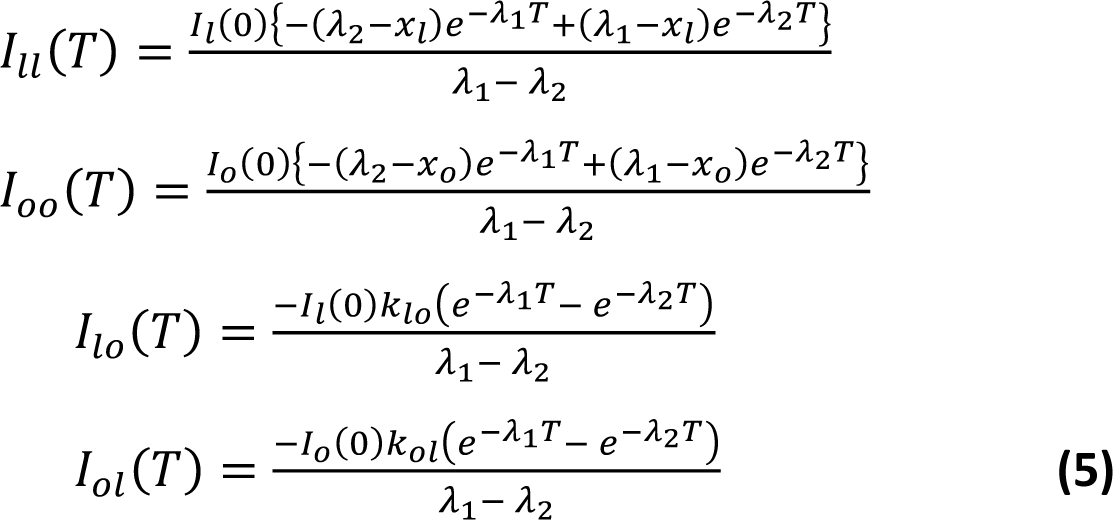

Where *λ*_1,2_, *x*_l_, and *x*_o_ are defined according to the following relationships:

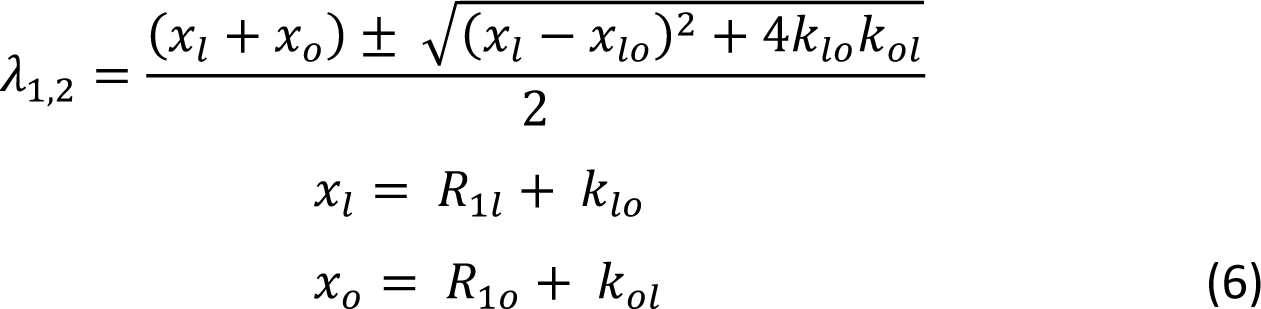

and the rate of transition from the locked to the open conformations, *k_lo_*, the rate of transition *vice versa*, *k*_ol_, the longitudinal relaxation rate in the locked conformation, *R*_1l_, as well as the longitudinal relaxation rate of the open conformation, *R*_1o_, were fitted. *I*_l_(0) and *I*_o_(0) denote the initial amounts of longitudinal carbon magnetization associated with the locked and the open conformations at the beginning of the mixing periods.

### NMR Diffusion

Translational diffusion coefficients were measured by recording a series of 1D ^13^C-edited spectra at different temperatures, using a pulse scheme (^13^C-edited BPP-LED) that is similar to a ^15^N-edited BPP-LED experiment with ^15^N and ^13^C pulses interchanged (*25*). To compensate for the different *J*-coupling of the methyl-groups compared to the amide groups, a gradient duration d of 3.2 ms was used instead of 4.8 ms used in the ^15^N-filtered version. In addition, a diffusion delay *T* of 400 ms and a *τ* of 0.1 ms were used. The strength of the encoding/decoding was increased stepwise. Transverse gradients instead of z-gradients were employed to minimize convection effects (*71*). The resulting ^1^H signal was integrated over the methyl ^1^H frequency range to obtain intensities as a function of encoding/decoding gradient strength. To adjust for the temperature dependent changes in viscosity it was adjusted based on published values for D_2_O-based buffers (*72*). In order to construct the calibration curve given in fig. S1J the following published translational diffusion coefficients, measured at 25°C, were used: Immunoglobin binding domain of streptococcal protein G, GB1 (MW = 6.2 kDa) (*73*), lysozyme (MW = 14.1 kDa) and Interleukin-10 (MW = 37.2 kDa) (*74*), bacterial HsIV (MW = 230 kDa), ½ proteasome from *T. acidophilum*, *α*7*α*7 (MW = 360 kDa), and the *α*7 single ring variant of the proteasome (MW = 180 KDa) (*75*).

### Bio-Layer Interferometry (BLI)

BLI experiments were performed on an Octet RED96 system (Fortébio) at 25°C as outlined in detail before (*76*). Briefly, the respective ligands were biotinylated using the biotinylation kit EZ-Link NHS-PEG4-Biotin (Thermo Fisher Scientific). The biotin label was resolved in H_2_O and directly added to the protein solution in a final molar ratio of 1:1 in 25 mM MES, pH 6.0 to ensure preferential labeling of the amino-terminal *α*-amino group, followed by gentle mixing at room temperature for 45 min. Unreacted biotin was removed with Zeba Spin Desalting Columns (7 MWCO, Thermo Fisher Scientific). Biotin-labeled proteins were immobilized on the streptavidin (SA) biosensors (Fortébio) and the biosensors were subsequently blocked with EZ-Link Biocytin (Thermo Fisher Scientific). Analytes were diluted and applied in a dose-dependent manner to the biosensors immobilized with the biotinylated ligand. Bovine serum albumin (BSA) powder (Sigma-Aldrich) and Tween-20 were added to final concentrations of 0.1% and 0.05%, respectively, to avoid non-specific interactions. Parallel experiments were performed for reference sensors with no analyte bound and the signals were subsequently subtracted during data analysis. The association period was set to 30 s and dissociation to 40 s. Data measurements and analysis were performed by using the Data acquisition 10.0 and the Data analysis HT 10.0 (Fortébio) software, respectively.

### Proteolytic cleavage assay

The fluorescent reporter peptides Abz-KASPVSLGY^NO2^D and p23 (Abz-GNWVSAAKFETYR^NO3^SKNTQDYGILQI), originally described in (*43*) and (*23*), respectively, were purchased from GenScript and prepared by dissolving in the lyophilized peptides directly in PBS (p23) or DMSO (Abz-KASPVSLGY^NO2^D). Cleavage assays were monitored using a FLUOstar Optima plate reader (BMG LABTECH), using an excitation wavelength of 320 nm and an emission wavelength of 440 nm, respectively, in 96-well black clear bottom microplates (Corning®). All experiments were prepared in PBS and contained either 0.2 μM of the respective DegP enzyme (monomer concentration) and 7.5 μM of p23 peptide or 10 μM of the enzyme with 50 μM β-Casein (Sigma-Aldrich) and 100 μM of the Abz-KASPVSLGY^NO2^D reporter peptide. All assays were performed as triplicates.

β-Casein degradation was, in addition, assessed *via* SDS-PAGE by incubating 1 μM DegP with 5 μM of β-Casein in PBS at 25 °C or 37 °C. At specific time intervals of 0 min, 5 min, 10 min, 20 min, 30 min, 60 min, and 90 min, respectively. Samples were taken out, mixed with sample loading dye, and subsequently heat-denatured at 95 °C for 10 minutes, before loading them onto SDS-PAGE gels. The samples were subsequently analyzed by SDS-PAGE using 4–20% Mini-Protean TGX–SDS-Gels (Bio-Rad).

## Acknowledgments

We thank G. Mas (Basel), P. Schanda (Grenoble), and R. Sprangers (Regensburg) for providing NMR pulse sequences. The Swedish NMR Centre of the University of Gothenburg is acknowledged for spectrometer time. B.M.B. gratefully acknowledges funding from the Wallenberg Centre for Molecular and Translational Medicine, University of Gothenburg, Sweden.

## Author contributions

B.M.B. conceived the study and designed the experiments together with D.Š.. D.Š. performed all experimental work. D.Š., J.T., and B.M.B. analyzed and discussed the data. D.Š., J.T., and B.M.B. wrote jointly the paper.

## Data and materials availability

All data needed to evaluate the conclusions in the paper are present in the paper and/or Supplementary Materials. The sequence-specific NMR resonance assignments for the PDZ1–PDZ2 (25°C and 50°C, respectively), PDZ1, and PDZ2-constructs were deposited in the BMRB under accession codes 50450–50453.

## Supplementary Materials

**Fig. S1.**
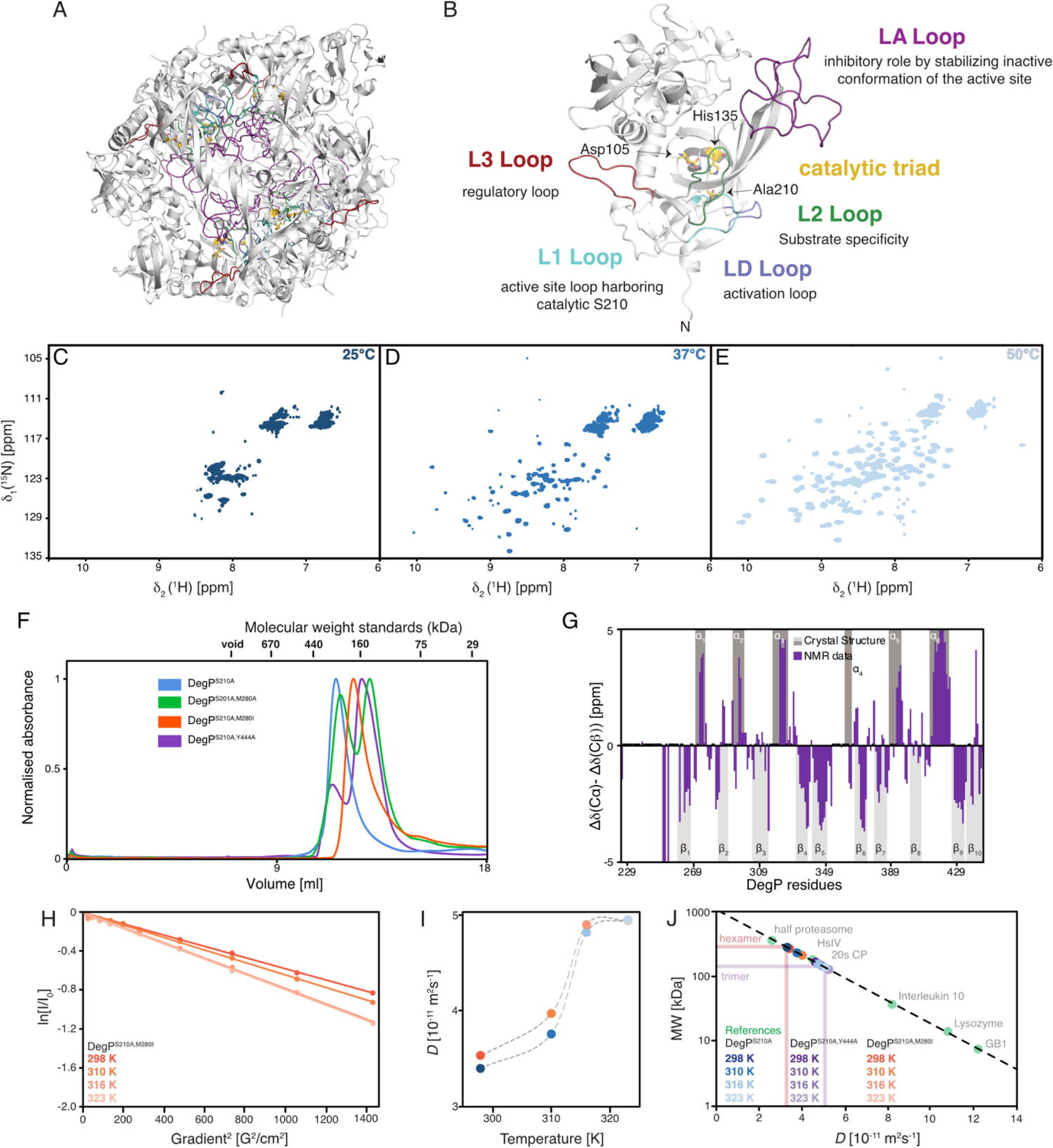
Solution studies of full-length DegP. (**A**) Structural model of the DegP^S210A^ in its inactive hexameric form highlighting the different regulatory loops involved in its proteolytic function. Based on the structural model developed by Figaj *et al.* (*18*). (**B**) Zoom on a DegP^S210A^ monomer with the highlighted regulatory loops together with the catalytic triad and their respective function. (**C–E**) 2D [^15^N,^1^H]-NMR spectrum of [*U*-^2^H,^15^N]-DegP^S210A^ measured at the indicated temperatures. (**F**) Gel elution profiles of different full-length DegP^S210A^ variants as indicated. Recorded at 8°C in NMR-buffer on a Superdex S200increase 10/300 column (GE Healthcare). The column void volume and the molecular weights of a standard calibration curve are indicated. (**G**) Secondary backbone ^13^C chemical shifts in solution of the full-length DegP^S210A,Y444A^ protein variant compared to the hexameric X-ray structure (PDB-ID: 1KY9). Positive values indicate α-helix whereas negative values β-sheet secondary structure elements. (**H**) Measurement of the molecular diffusion constants with a ^13^C-methyl-filtered diffusion experiment for DegP^S210A,M280I^. The logarithm of the signal intensity is plotted against the squared gradient strength of the applied pulsed-field gradients. [*U*-^2^H, Ile-*δ*_1_-^13^CH_3_]–DegP^S210A,M280I^ was measured at different temperatures as indicated. Solid lines are a linear fit to the data. (**I**) Obtained molecular diffusion constants for DegP^S210A^ (blue; data also shown in Fig. 1F) and DegP^S210A,M280I^ (orange) plotted against the temperature. The broken lines serve as a guide to the eyes only. (**J**) Calibration curve for estimating the molecular weight based on the determined molecular diffusion constants. Linear correlation of the molecular weight (MW: logarithmic scale) plotted against the molecular diffusion constants. Published diffusion constants of proteins with known molecular weight are indicated in green (see Methods for details). Temperature dependent data for DegP^S210A^, DegP^S210A,Y444A^, and DegP^S210A,M280I^ are shown as blue, purple or orange gradients, respectively, as indicated.

**Fig. S2.**
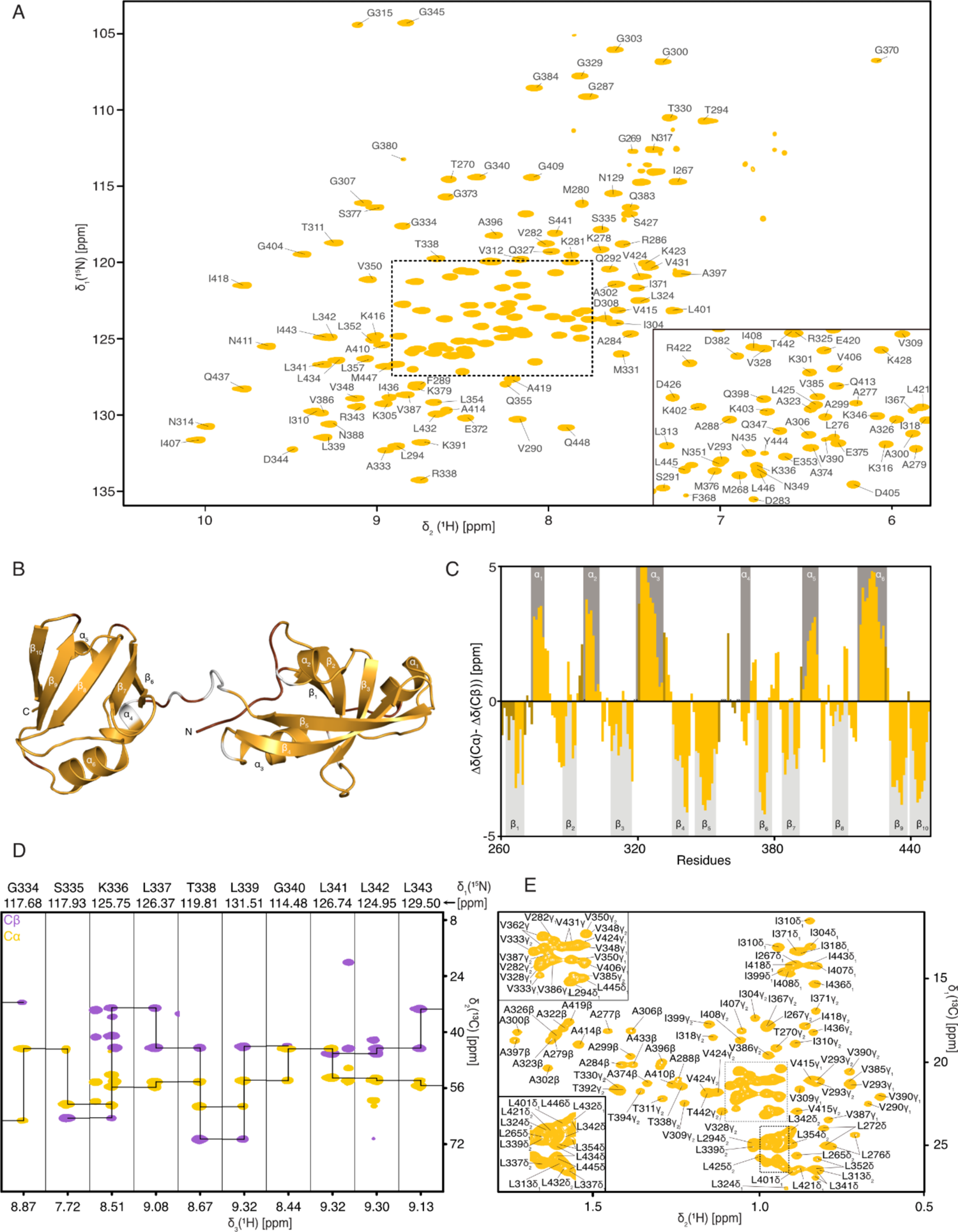
The isolated PDZ1–PDZ2 domain construct in solution. (**A**) 2D [^15^N,^1^H]-NMR-spectrum of [*U*-^15^N,^13^C]-PDZ1–PDZ2. The sequence-specific resonance assignment obtained from 3D triple-resonance experiments is indicated. (**B**) Cartoon representation of the crystal structure of the PDZ1–PDZ2 domains (PDB-ID: 3OU0) with the assigned resonances highlighted in orange and unassigned residues in white. (**C**) Secondary backbone ^13^C chemical shift analysis plotted against the PDZ1–PDZ2 residues. Dark shaded bars present residues experiencing line-broadening in [^15^N,^1^H]-NMR-spectra that could be assigned unambiguously *via* 2D [^13^C,^1^H]-NMR-spectra based TOCSY-type 3D experiments. The secondary structure elements derived from the crystal structure (PDB-ID: 3OU0) are highlighted in grey. (**D**) Representative strips for residues G74–L83 from a 3D HNCACB spectrum of the PDZ1– PDZ2-domains. (**E**) Methyl-region of a [^13^C,^1^H]-NMR-spectrum of [*U*-^15^N,^13^C]-PDZ1–PDZ2 with the assignment of the methyl side-chain resonances indicated.

**Fig. S3.**
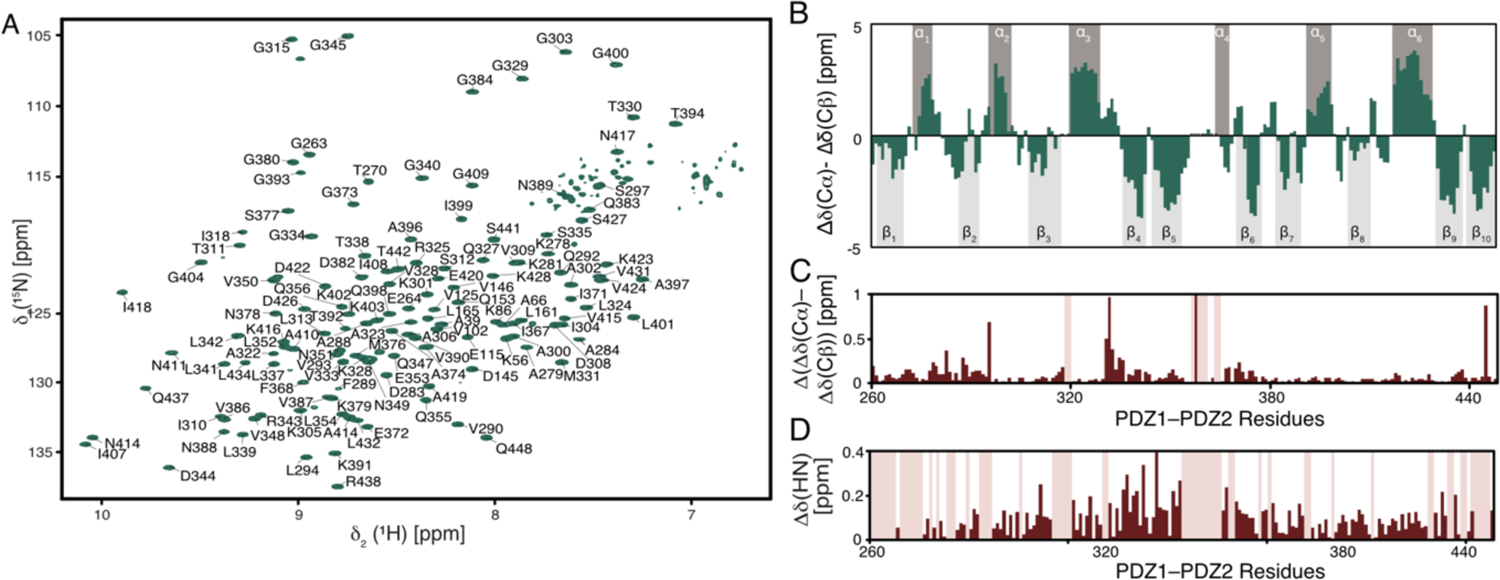
The isolated PDZ1–PDZ2 domain construct in solution at low temperature. (**A**) 2D [^15^N,^1^H]-NMR-spectrum of [*U*-^15^N,^13^C]-PDZ1–PDZ2 at 25°C. The sequence-specific resonance assignment obtained from 3D triple-resonance experiments is indicated. (**B**) Secondary backbone ^13^C chemical shift analysis plotted against the PDZ1–PDZ2 residues. The secondary structure elements derived from the crystal structure (PDB-ID: 3OU0) are highlighted in grey. (**C**) The difference of the secondary backbone ^13^C chemical shifts at the different temperatures (*Δ*(*Δδ*(^13^C*α*)-*Δδ*(^13^C*β*)) are shown in dark red. (**D**) Combined chemical shift differences of the amide moieties plotted versus the PDZ1–PDZ2 amino acid residue number. In panels, **C** and **D** broadened residues are indicated by the red shading.

**Fig. S4.**
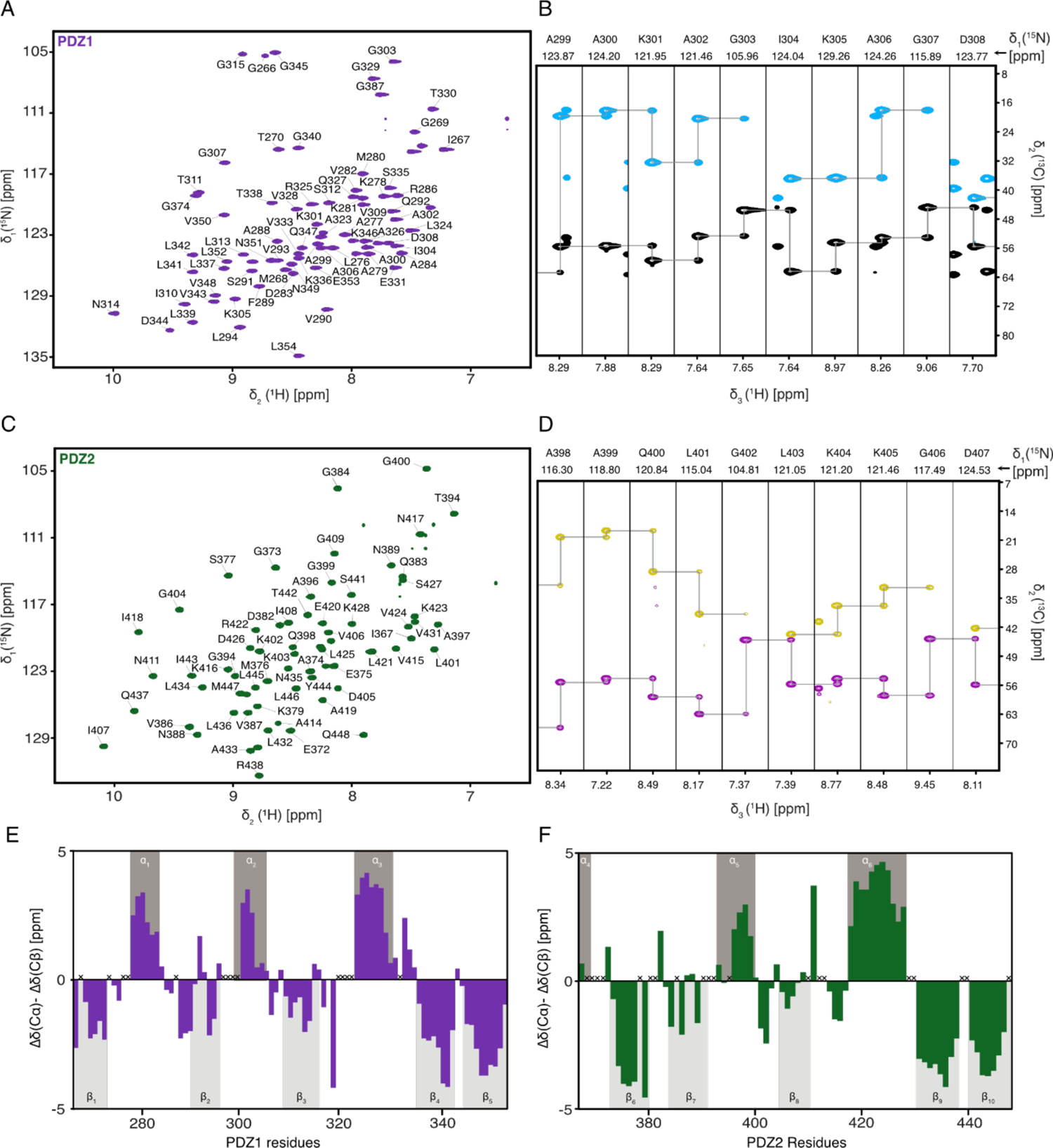
The isolated PDZ1 and PDZ2 domains in solution. (**A**) 2D [^15^N,^1^H]-NMR-spectrum of [*U*-^15^N,^13^C]-PDZ1. The sequence-specific resonance assignment obtained from 3D triple-resonance experiments is indicated. (**B**) Representative strips for residues A299–D308 from a 3D HNCACB spectrum of the PDZ1-domain. (**C**) 2D [^15^N,^1^H]-NMR-spectrum of [*U*-^15^N,^13^C]-PDZ1. The sequence-specific resonance assignment obtained from 3D triple-resonance experiments is indicated. (**D**) Representative strips for residues A398–D407 from a 3D HNCACB spectrum of the PDZ1-domain. (**E, F**) Secondary backbone ^13^C chemical shift analysis plotted against the PDZ1 (**E**) and PDZ2 (**F**) residues. The secondary structure elements derived from the crystal structure are highlighted in grey.

**Fig. S5.**
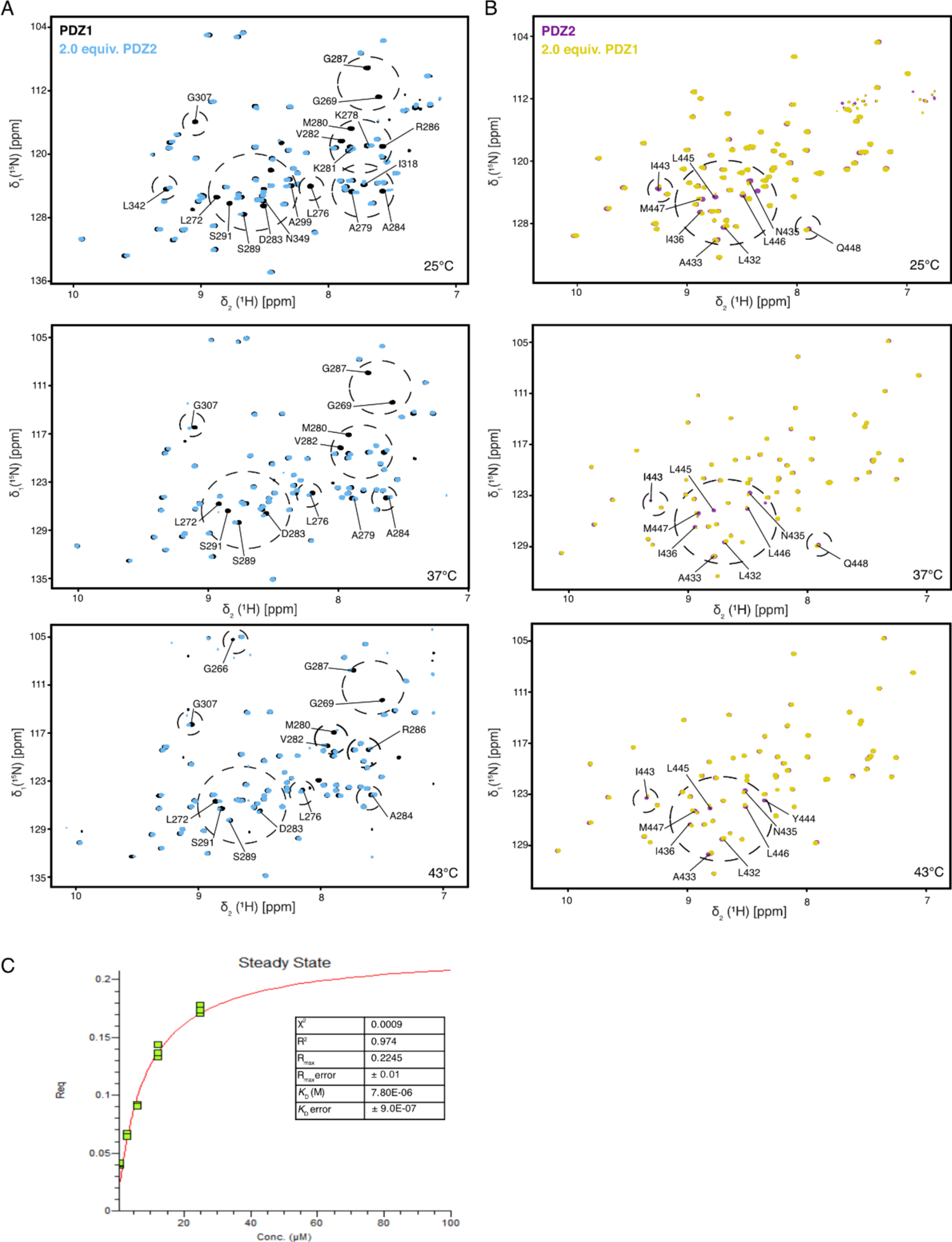
Temperature dependence of the PDZ1:PDZ2 interaction. (**A, B**) 2D [^15^N,^1^H]-NMR spectra of [*U*-^15^N]-PDZ1 (**A**, purple) and [*U*-^15^N]-PDZ2 (**B**, black) as well as after the addition of two molar equivalents of the respective other domain. Titrations were measured at 25°C, 37°C, and 43°C, as indicated. (**C**) Steady-state analysis of the BLI-curves shown in Fig. 2F. Curves were measured and analyzed in triplicates (green data points). The red curve is the least-squares fit to the data. The resulting *K*_D_ and indicators for the fitting quality (R^2^,*χ*^2^) are indicated.

**Fig. S6.**
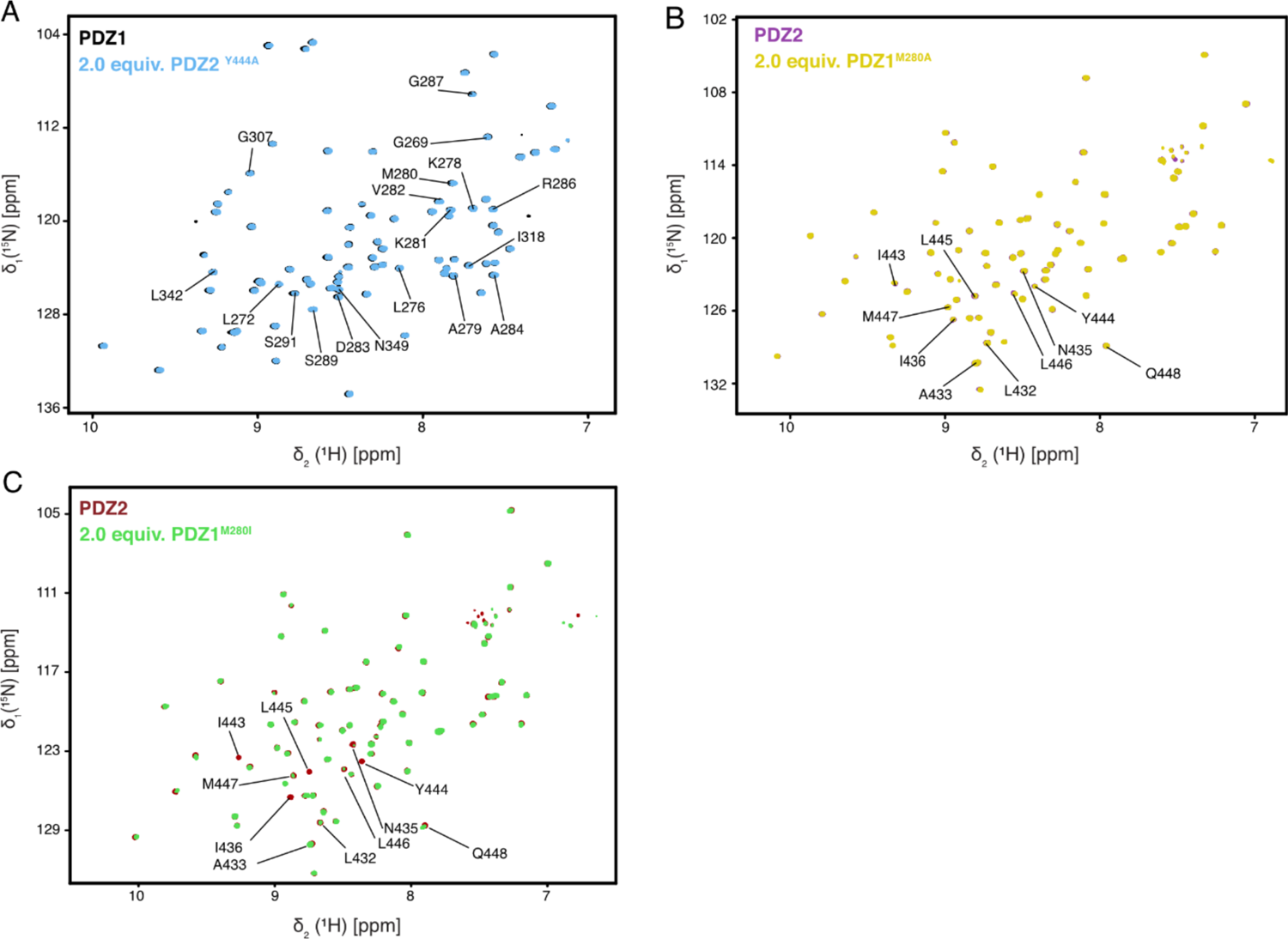
Interactions of the PDZ1^M280A^, PDZ1^M280I^, and PDZ2^Y444A^ mutants. (**A, B, C**), 2D [^15^N,^1^H]-NMR spectra of [*U*-^15^N]-PDZ1 (**A**, black), [*U*-^15^N]-PDZ2 (**B**, black, **C**, red) as well as after addition of two molar equivalents of the respective other domain carrying the indicated single-point mutation. Titrations were measured at 25°C.

**Fig. S7.**
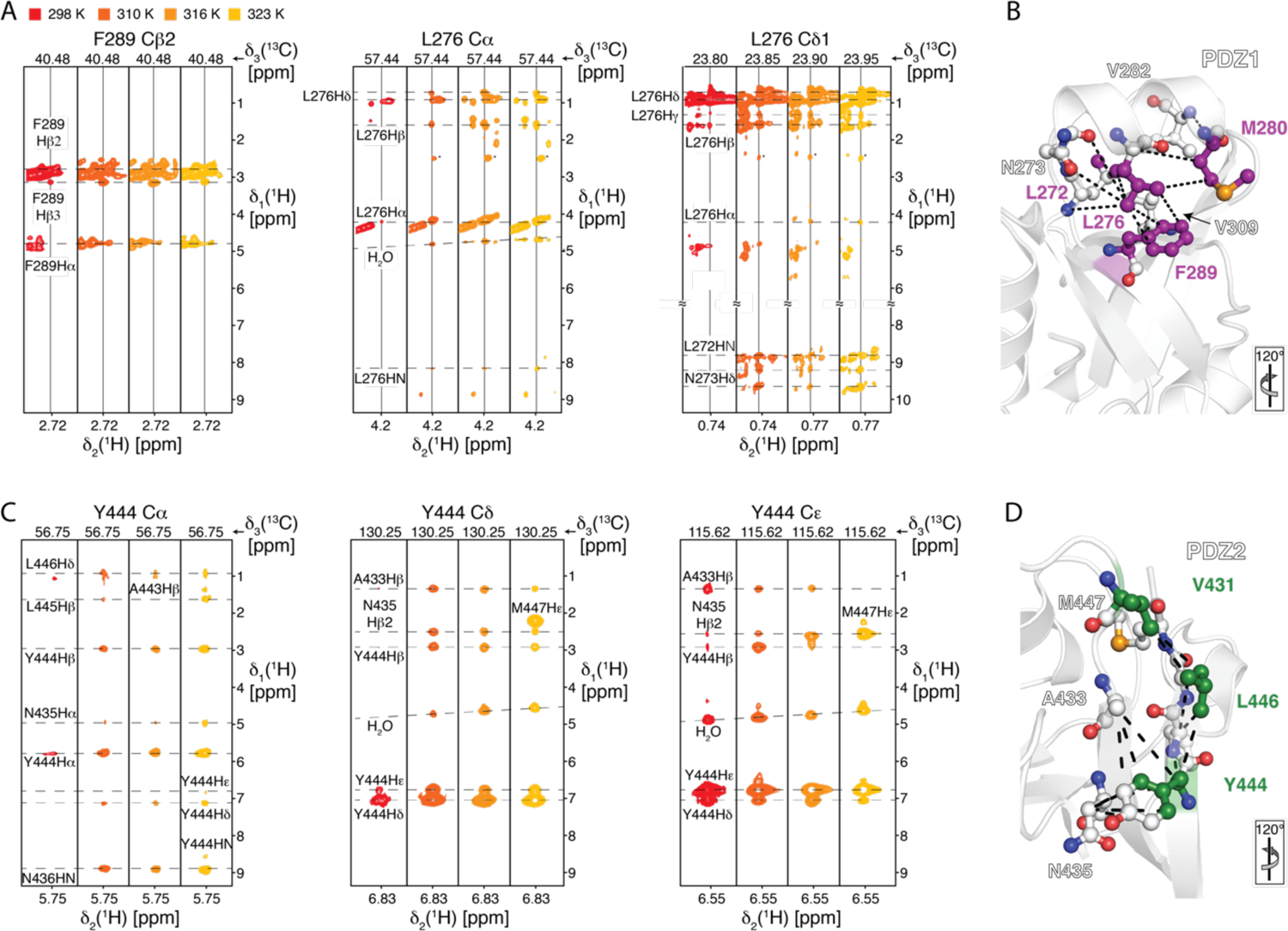
Temperature-dependent analysis of the NOE-network stabilizing the PDZ1–PDZ2 interface. (**A**) Representative NOE strips from NOESY experiments at the indicated temperatures. Intramolecular NOE cross-peaks showing parts of the NOE-network on PDZ1 stabilizing the interaction regions around M280 is indicated. (**B**) Visualization of the detected intramolecular NOEs (dashed lines) on the structure of PDZ1. (**C**) Representative NOE strips from NOESY experiments at the indicated temperatures. Intramolecular NOE cross-peaks showing parts of the NOE-network on PDZ2 stabilizing the interaction regions around Y444 is indicated. (**D**) Visualization of the detected intramolecular NOEs (dashed lines) on the structure of PDZ2. (**B, D**) Broadened residues upon domain interaction at low temperature are indicated in purple (PDZ1) and green (PDZ2) as shown in Fig. 3H as well as the respective orientation.

**Fig. S8.**
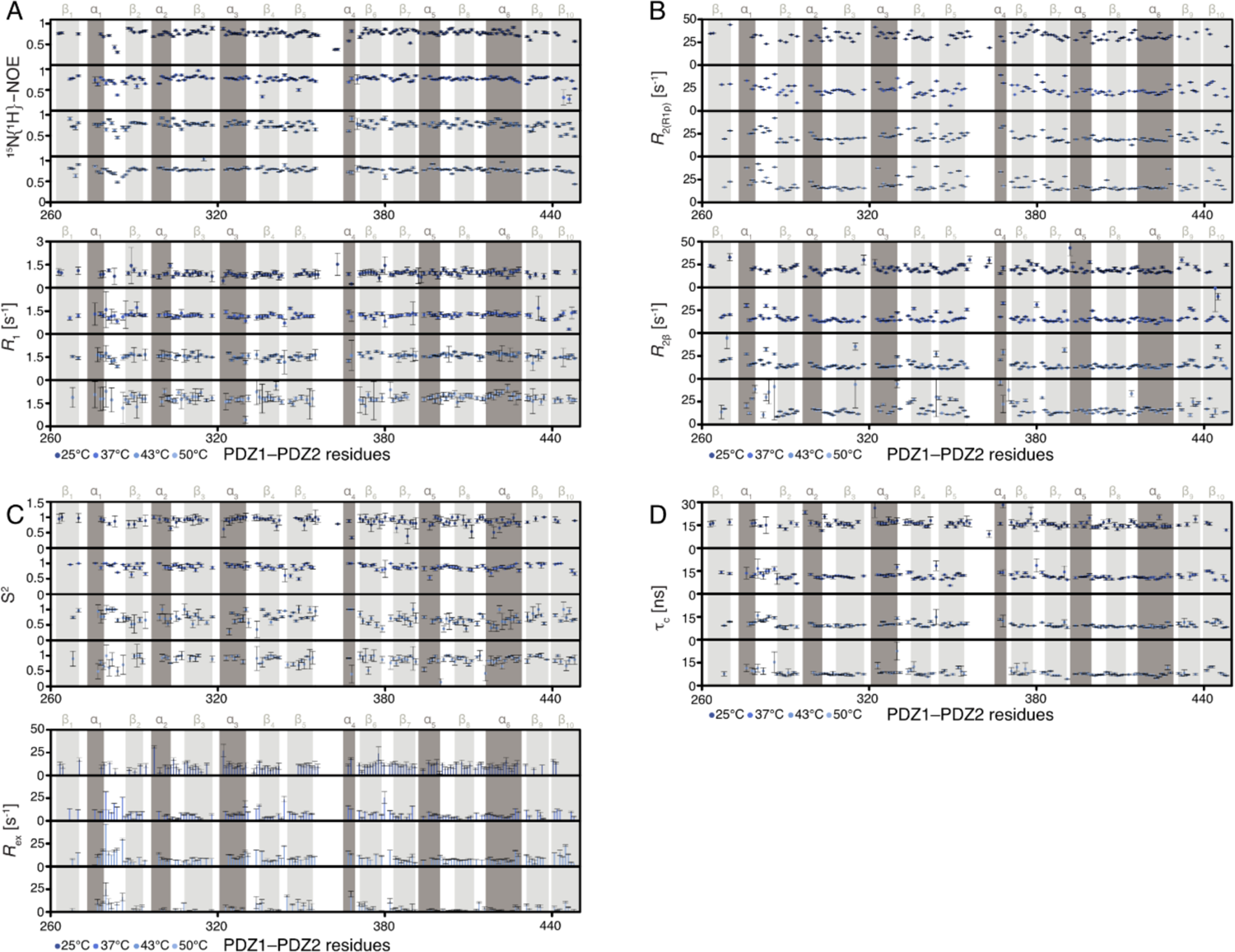
Temperature dependence of the PDZ1-PDZ2 backbone dynamics. (**A**) Local backbone dynamics on the pico– to nanosecond timescale probed by hetNOE and *R*_1_ measurements. (**B**) Obtained transversal relaxation rates (*R*_2β_ and *R*_2(R1ρ)_) reporting on micro- to millisecond motions. (**C**) Temperature dependence of the sub-nanosecond order parameter S^2^ and the exchange rate *R*_ex_, reporting on possible conformational exchange contributions on the micro- to millisecond timescale, calculated using the Lipari-Szabo model-free approach. (**D**) Rotational correlation time (*τ*_c_) is determined from the ratio of *R*_1_ and *R*_2(R1ρ)_. All data are plotted against the PDZ1–PDZ2 residues at the indicated temperatures ranging from 25–50°C.

**Fig. S9.**
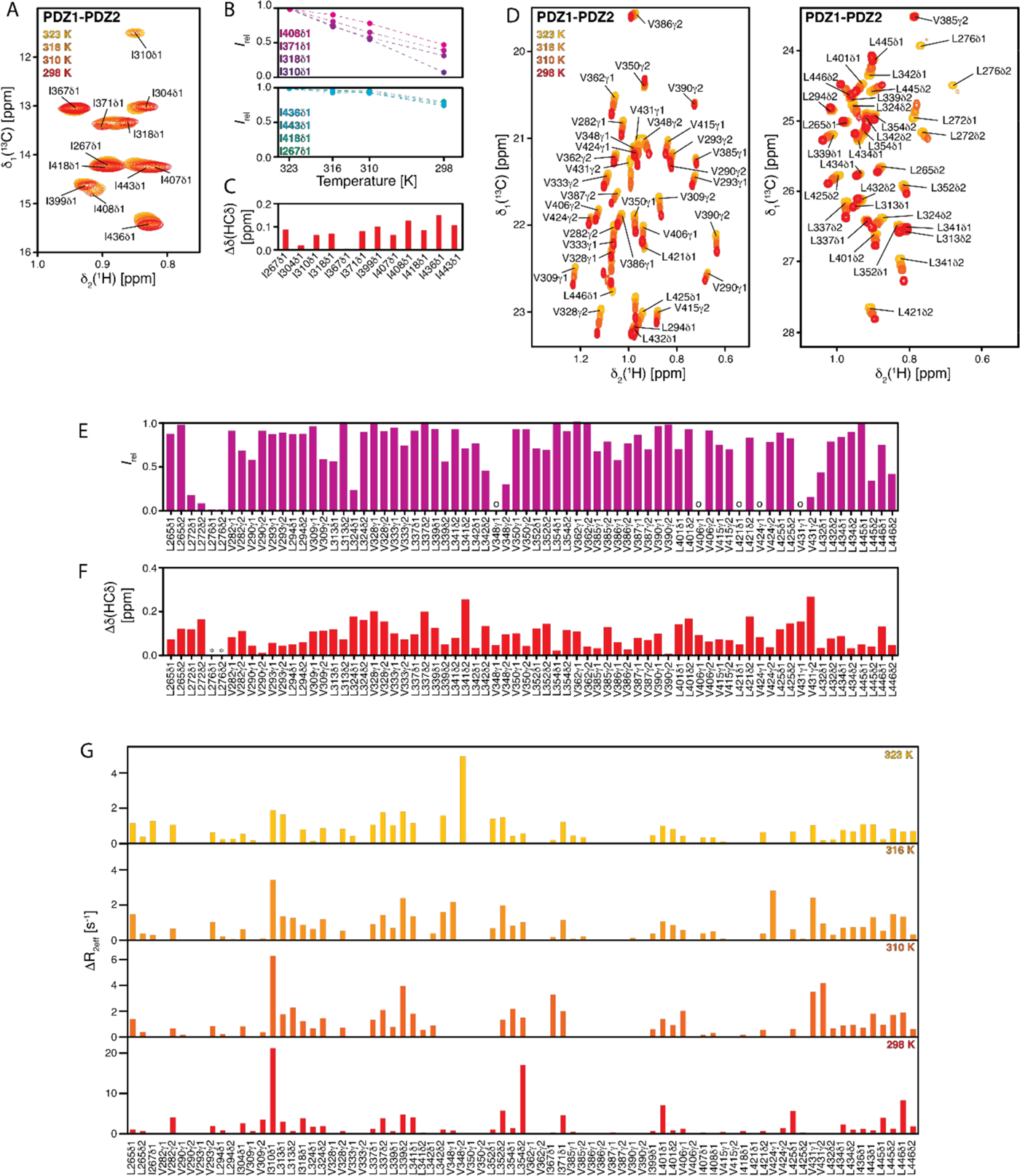
Methyl group dynamics. (**A**) Isoleucine methyl region of a 2D [^13^C,^1^H]-NMR spectrum of [*U*-^13^C,^15^N]-PDZ1–PDZ2 at different temperatures as indicated. (**B, C**) The temperature dependence of the isoleucine signal intensities (**B**) and chemical shift changes (**C**) for the PDZ1–PDZ2 construct. The broken lines in panel **B** serve as guide to the eyes only. (**D**) Leucine and valine methyl region of a 2D [^13^C,^1^H]-NMR spectrum of [*U-*^2^H, Leu-δ_1,2_-Val-γ_1,2_^13^CH_3_]*-*PDZ1–PDZ2 at different temperatures as indicated. (**E, F**) The temperature dependence of the leucine/valine signal intensities (**E**) and chemical shift changes (**F**) for the PDZ1–PDZ2 construct. o in panel **E** indicates overlapping peaks, whereas * in panel **F** indicates severe line-broadening beyond detection. (**G**) Δ*R*_2eff_ values for isoleucine, leucine, and valine methyl groups, obtained from the difference of *R*_2eff_ at the lowest and highest CPMG frequency υ_CPMG_, at the indicated temperatures.

**Fig. S10.**
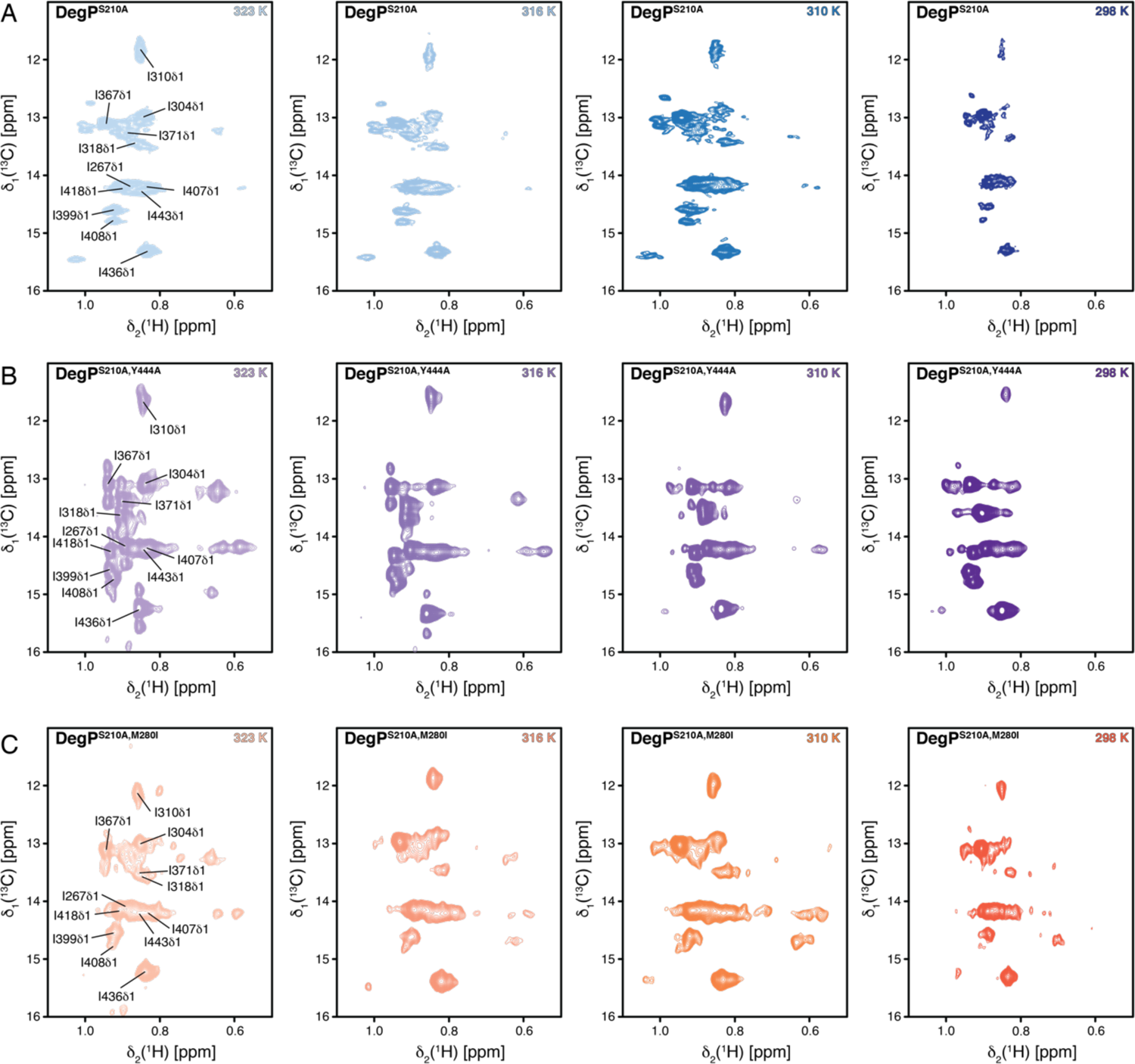
Isoleucine spectra of full-length DegP^S210A^-variants. (**A–C**) 2D [^13^C,^1^H]-NMR spectra of [*U-*^2^H, Ile-δ_1_-^13^CH_3_]*–*DegP^S210A^ (**A**), [*U-*^2^H, Ile-δ_1_-^13^CH_3_]*–*DegP^S210A,Y444A^ (**B**), and [*U-*^2^H, Ile-δ_1_-^13^CH_3_]*–*DegP^S210A,M280I^ (**C**) measured at the indicated temperatures. The sequence-specific resonance assignment of selected isoleucine resonances derived from the isolated PDZ1–PDZ2 domain are indicated.

**Fig. S11.**
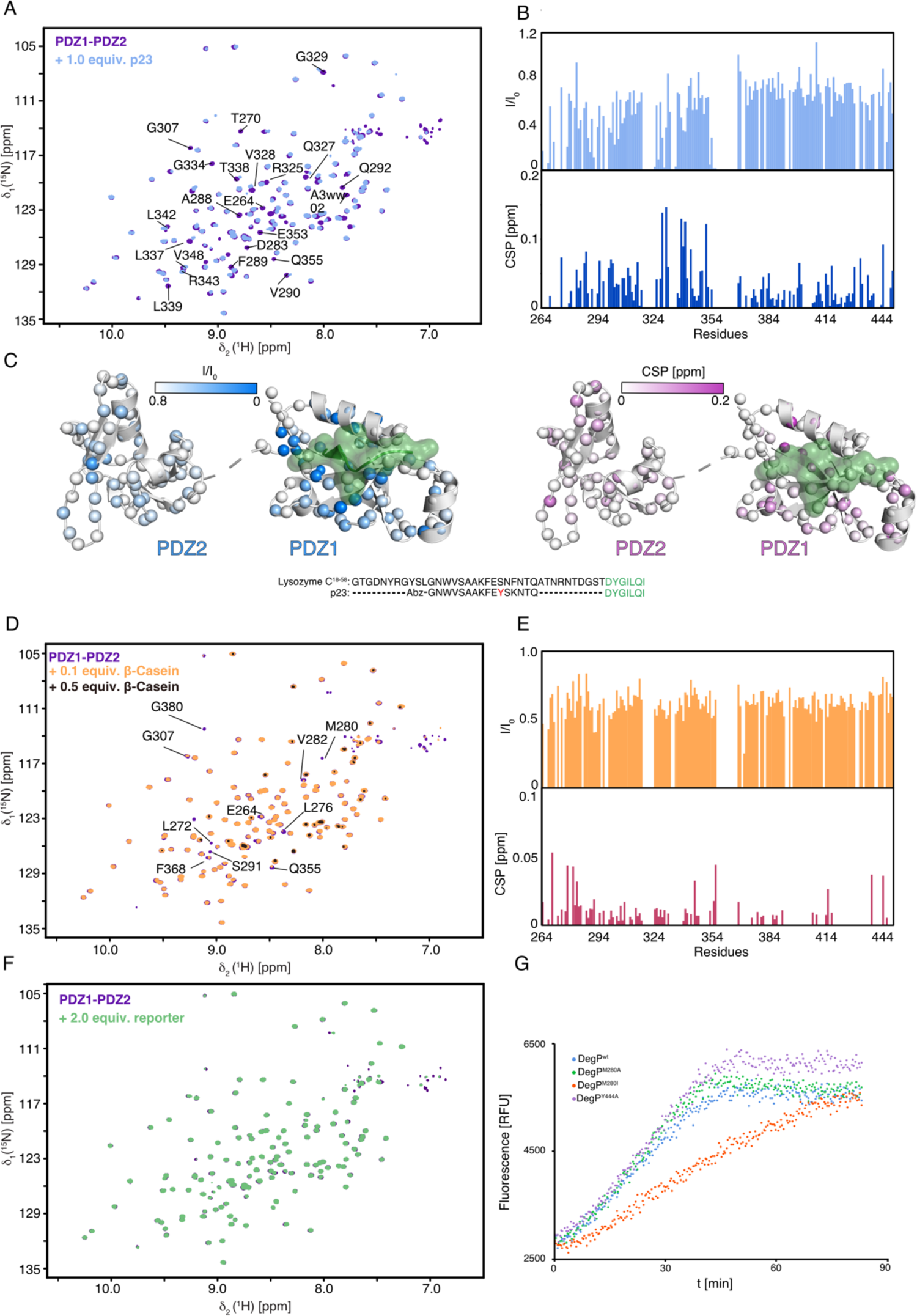
PDZ1-PDZ2 construct interactions with different substrates. (**A**) 2D [^15^N,^1^H]-NMR spectra of [*U*-^15^N]-PDZ1–PDZ2 (purple) and after the addition of 1 molar equivalent of p23 (blue). All NMR titrations were measured at 37°C. (**B**) Residue-resolved backbone amide NMR signal attenuation (I/I_0_) and detected signal chemical shift perturbations (CSP) of the p23 titration. (**C**) Observed p23 titration effects plotted on the previously reported crystal structure of the DegP:Lysozyme C^15-58^ (green) complex (3OTP) (*14*). The amide moieties of the PDZ1–PDZ2 construct are shown as spheres. Signal attenuation values are indicated by the white to blue and CSP values by the white to the purple gradient. Inset: Sequence alignment of Lysozyme C^15–58^ and p23 for the visible residues highlighted in green. (**D**) 2D [^15^N,^1^H]-NMR spectra of [*U*-^15^N]-PDZ1–PDZ2 (purple) and after the addition 0.1 (orange) and 0.5 (black) molar equivalents of β-Casein. (**E**) Residue-resolved backbone amide NMR signal attenuation (I/I_0_) and detected signal chemical shift perturbations (CSP) of the β-Casein titration (0.1 equivalents). (**F**) 2D [^15^N,^1^H]-NMR spectra of [*U*-^15^N]-PDZ1–PDZ2 (purple) and after the addition 2 molar equivalents of reporter peptide. (**G**) Proteolysis of p23 (7.5 µM) by DegP and its variants (0.2 µM of monomer concentration) at 25°C.

**Fig. S12.**
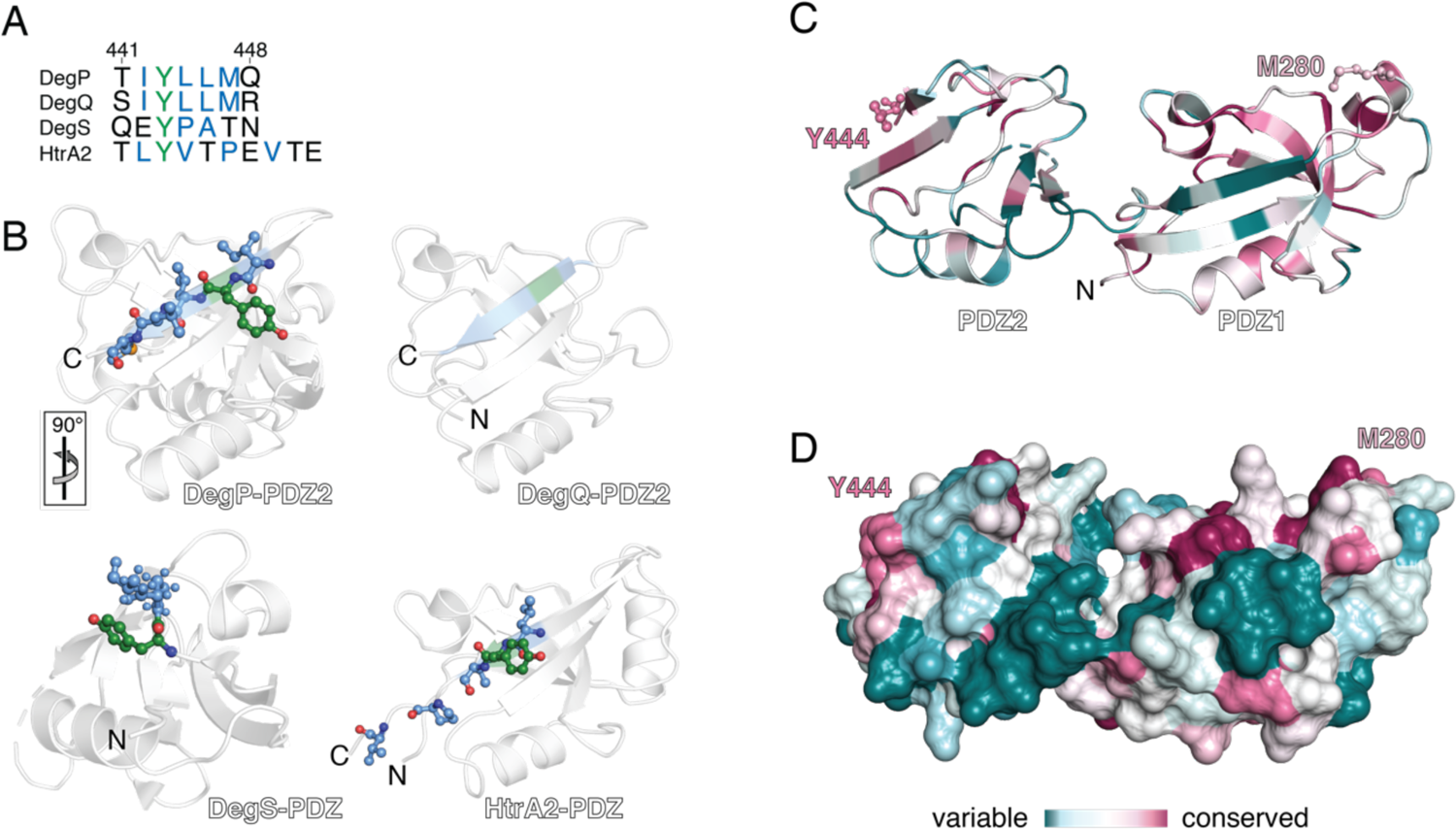
Sequence conservation analysis of the interdomain lock. (**A**) Sequence comparison of the *β*_10_-strand containing Y444 in the PDZ of DegP with corresponding domains of related bacterial periplasmic DegQ and DegS as well as human mitochondrial HtrA2. (**B**) Cartoon representation of the DegP-PDZ2 (PDB-ID: 3OU0) with focus on Y444 and adjacent hydrophobic residues forming a hydrophobic patch facilitating the inter-domain lock (*23*). For the DegQ-PDZ2 only a medium resolution structure at 7.5Å (PDB-ID: 4A8C) is available without any side-chain information (*22*). In contrast to DegP and DegQ, DegS has only a single PDZ-domain (PDB-ID: 4RQY). Similarly, human mitochondrial HtrA2 bears also a single PDZ-domain (PDB-ID: 1LCY), for which recently the involvement of the highlighted tyrosine in oligomeric resting state stabilization could be shown (*21*). (**C, D**) Cartoon **(C)** and surface **(D)** representation of the DegP PDZ1-PDZ2 domains displaying the degree of conservation with the indicated gradient ranging from 0 (variable) to 9 (conserved). Positions of M280, 7/9 conservation score, and Y444, 8/9 conservation score are indicated. Analysis was performed *via* the ConSurf (Tel Aviv University, Israel) server (*78, 79*).

**Table S1.** Summary of the obtained diffusion coefficients

| Temperature | DegP<sup>S210A</sup> | DegP<sup>S210A,Y444A</sup> | DegP<sup>S210A,M280I</sup> |
| --- | --- | --- | --- |
| 25°C | 3.41•10<sup>-7</sup> cm<sup>2</sup>s<sup>-1</sup> | 4.64•10<sup>-7</sup> cm<sup>2</sup>s<sup>-1</sup> | 3.53•10<sup>-7</sup> cm<sup>2</sup>s<sup>-1</sup> |
| 37°C | 3.81•10<sup>-7</sup> cm<sup>2</sup>s<sup>-1</sup> | 4.81•10<sup>-7</sup> cm<sup>2</sup>s<sup>-1</sup> | 3.97•10<sup>-7</sup> cm<sup>2</sup>s<sup>-1</sup> |
| 43°C | 4.83•10<sup>-7</sup> cm<sup>2</sup>s<sup>-1</sup> | 4.96•10<sup>-7</sup> cm<sup>2</sup>s<sup>-1</sup> | 4.89•10<sup>-7</sup> cm<sup>2</sup>s<sup>-1</sup> |
| 50°C | 4.94•10<sup>-7</sup> cm<sup>2</sup>s<sup>-1</sup> | 4.99•10<sup>-7</sup> cm<sup>2</sup>s<sup>-1</sup> | 4.94•10<sup>-7</sup> cm<sup>2</sup>s<sup>-1</sup> |

**Table S2.**
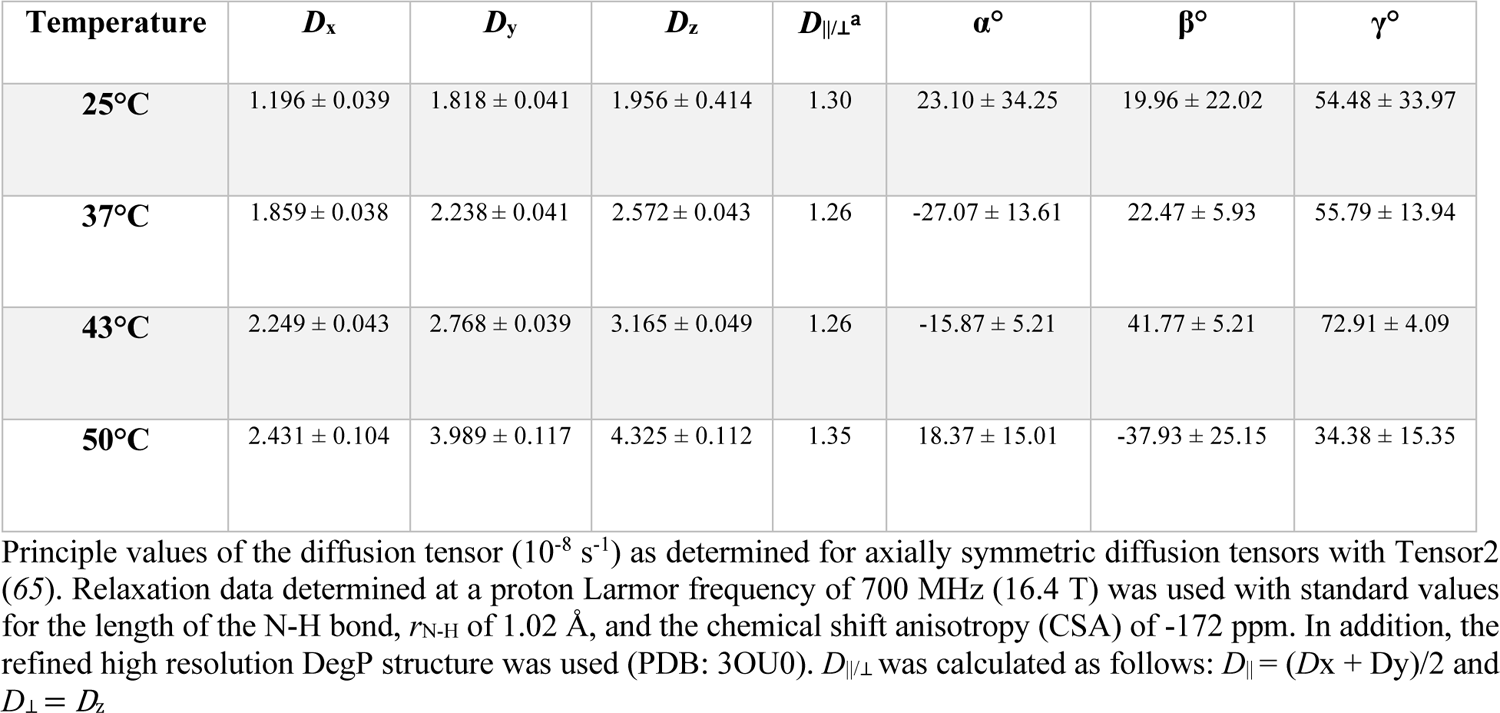
Summary of the calculated diffusion tensor values

**Table S3.**
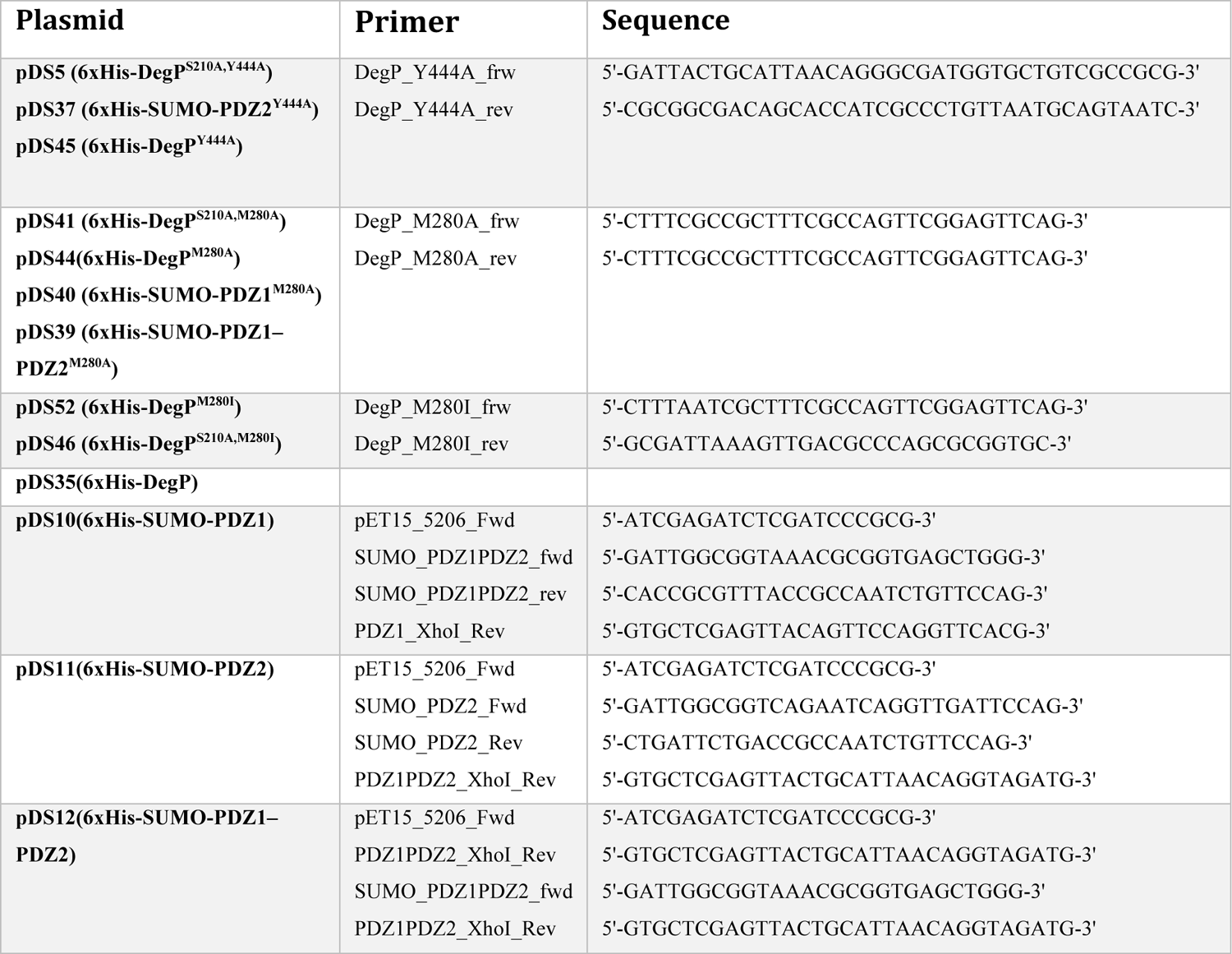
Plasmids and respective primers used in this study.

